# Engineering a seven enzyme biotransformation using mathematical modelling and characterized enzyme parts

**DOI:** 10.1101/603795

**Authors:** William Finnigan, Rhys Cutlan, Radka Snajdrova, Joseph P. Adams, Jennifer A. Littlechild, Nicholas J. Harmer

**Affiliations:** Department of Biosciences, Henry Wellcome Building for Biocatalysis, Stocker Road, Exeter EX4 4QD, U.K.; Living Systems Institute, Stocker Road, Exeter EX4 4QD, U.K.; GlaxoSmithKline R&D Ltd., Medicines Research Centre, Gunnels Wood Road, Stevenage, Hertfordshire SG1 2NY, U.K.

**Keywords:** Enzyme cascade, Kinetics, Cofactor recycling, Pathway optimization, Isolated enzymes, Reaction engineering, *in vitro* biocatalysis.

## Abstract

Multi-step enzyme reactions offer considerable cost and productivity benefits. Process models offer a route to understanding the complexity of these reactions, and allow for their optimization. Despite the increasing prevalence of multi-step biotransformations, there are few examples of process models for enzyme reactions. From a toolbox of characterized enzyme parts, we demonstrate the construction of a process model for a seven enzyme, three step biotransformation using isolated enzymes. Enzymes for cofactor regeneration were employed to make this *in vitro* reaction economical. Good modelling practice was critical in evaluating the impact of approximations and experimental error. We show that the use and validation of process models was instrumental in realizing and removing process bottlenecks, identifying divergent behavior, and for the optimization of the entire reaction using a genetic algorithm. We validated the optimized reaction to demonstrate that complex multi-step reactions with cofactor recycling involving at least seven enzymes can be reliably modelled and optimized.

**Significance statement:** This study examines the challenge of modeling and optimizing multi-enzyme cascades. We detail the development, testing and optimization of a deterministic model of a three enzyme cascade with four cofactor regeneration enzymes. Significantly, the model could be easily used to predict the optimal concentrations of each enzyme in order to get maximum flux through the cascade. This prediction was strongly validated experimentally. The success of our model demonstrates that robust models of systems of at least seven enzymes are readily achievable. We highlight the importance of following good modeling practice to evaluate model quality and limitations. Examining deviations from expected behavior provided additional insight into the model and enzymes. This work provides a template for developing larger deterministic models of enzyme cascades.

## Introduction

### Multi-step biocatalysis

Biocatalysis, the use of isolated enzymes or whole cells to perform chemical reactions, is increasingly the route of choice in the chemical and particularly the pharmaceutical industries (1). There is increasing interest in developing one-pot reactions, where multiple catalysts work together to complete chemical pathways. Multi-step pathways offer excellent cost and productivity benefits (2), driving reversible processes to completion without the need to isolate substrates or products at each step (3). Enzymes are well suited to these reactions, operating in similar aqueous conditions at low temperatures and pressures. In contrast, the use of several chemical steps together will often have differing operational requirements (4). Using isolated enzymes in biotransformations offers advantages (5) including reduced complexity and cost in downstream processing (6), fewer side reactions (7) and the removal of rate limiting diffusion across cellular membranes (8). Most attractively, parameters such as enzyme or substrate concentration, co-solvents, pH or temperature, can easily be manipulated (9). Furthermore, a mathematical model can be built and validated so that new pathways or components can be rapidly engineered and tested *in silico*.

The carboxylic acid reductases (CARs) are enzymes that have increasing interest as biocatalysts (10,11). CARs catalyze the reduction of carboxylic acids to aldehydes in mild conditions, and can connect many types of enzyme reaction allowing the construction of novel multi-enzyme pathways (Figure 1A) (12–14). Previously, we have shown CARs to be a fairly promiscuous enzyme class, catalyzing the reduction of both aliphatic and aromatic carboxylic acids. Electron rich acids are favored as the first step in the CAR reaction mechanism since attack by the carboxylate on the α-phosphate of ATP, is limiting (15). CARs have been shown to be useful in the construction of pathways for *in vivo* use. Some examples include the production of the flavor vanillin by yeast (16) and a synthetic pathway for the production of propane in *Escherichia coli* (17). The use of CARs as an isolated enzyme in a multistep reaction, however, has not been explored. Possibly this is due to the challenging and costly requirements for both ATP and NADPH regeneration (18), or the buildup of product inhibition by pyrophosphate (PP_I_) (15,19).

**Figure 1.**
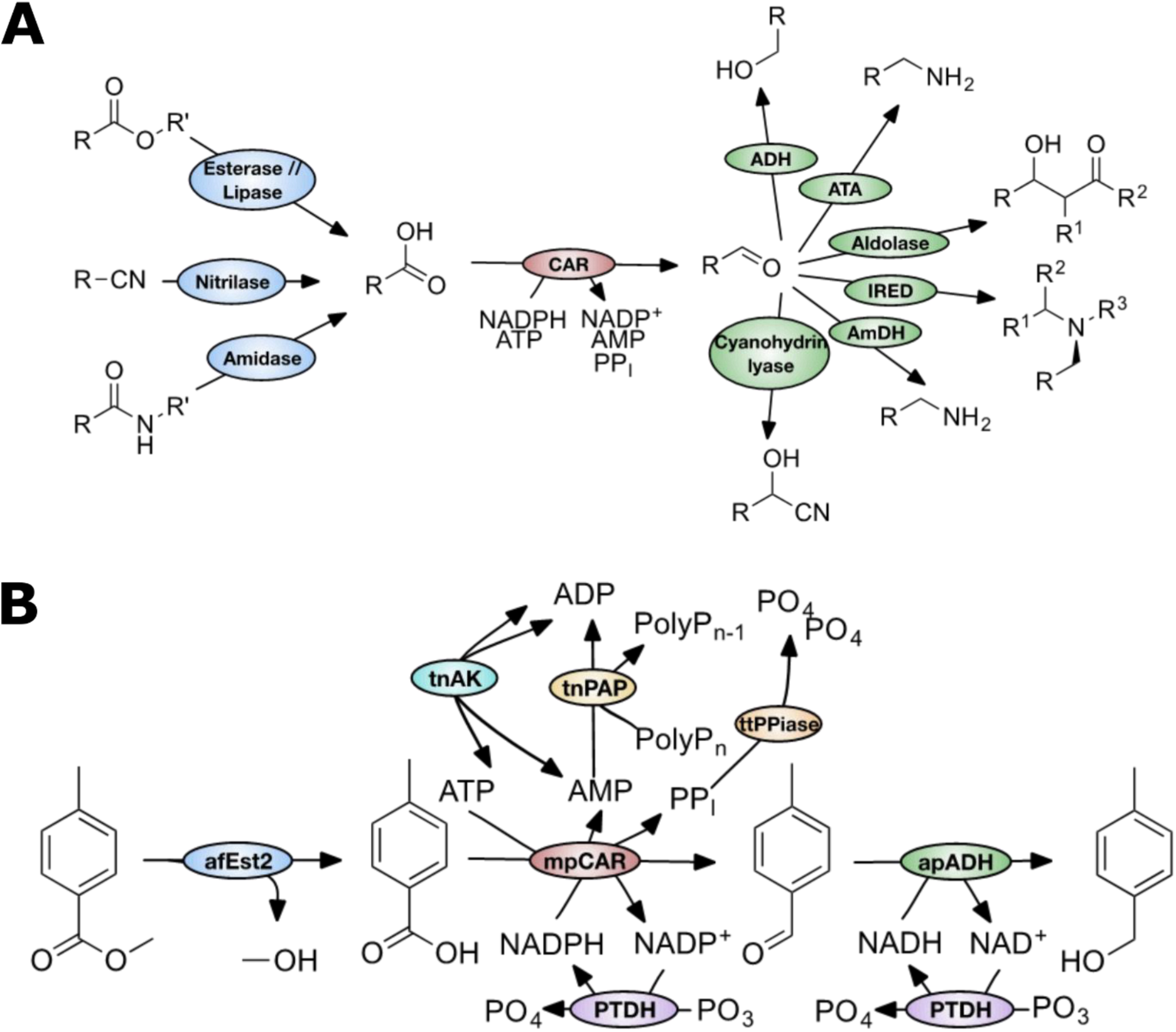
Utilizing CARs in multi-step enzyme reactions. A. CARs join many industrially relevant enzyme reactions, making them useful for the construction of novel multi-step enzyme reactions. Cofactors and additional substrates are not shown for enzymes other than CAR for clarity. ADH: alcohol dehydrogenase, ATA: amino transferase, IRED: imine reductase, AmDH: amine dehydrogenase, CAR: carboxylic acid reductase. R: R-group, limited by enzyme substrate specificity. B. A schematic of the seven enzyme reaction. The hydrolysis of methyl 4-toluate to 4-toluic acid, followed by reduction to 4-tolualdehyde and further to 4-tolylalcohol is shown. The use or production of water is not shown. afEst2: Esterase enzyme from *Archaeoglobus fulgidus*, mpCAR: Carboxylic acid reductase from *Mycobacterium phlei*, apADH: Alcohol dehydrogenase from *Aeropyrum pernix*, PTDH: Engineered phosphite dehydrogenase from *Pseudomonas stutzeri*, ttPPiase: Inorganic pyrophosphatase from *Thermus thermophilus*, tnPAP: Polyphosphate AMP phosphotransferase from *Thermodesulfobium narugense*, tnAK: Adenylate kinase from *Thermotoga neapolitana*. PolyP_n_: A polyphosphate molecule with a chain length of n phosphates.

### Cofactor regeneration

Many oxidoreductases require cofactors, most commonly the comparatively expensive NAD(P)H. For biocatalysis to be an economically viable process *in vitro*, cofactor regeneration is essential. Many enzyme systems have been developed for the regeneration of NAD(P)H (20). One such system is the phosphite dehydrogenase (PTDH) enzyme from *Pseudomonas stutzeri*. The wild-type enzyme regenerates NADH via the nearly irreversible oxidation of phosphite to phosphate. The enzyme has been engineered to regenerate NADPH and improve its thermostability (21,22), making it more attractive when regeneration of both NADH and NADPH is required.

Techniques for the regeneration of ATP are less well developed. Enzymes such as pyruvate kinase (PK), creatine kinase (CK), adenylate kinase (AK) and polyphosphate kinase (PPK) have been used (23). Of these, polyphosphate kinase makes use of the cheapest and most stable substrate, polyphosphate. Polyphosphate is also a substrate for polyphosphate-AMP phosphotransferase (PAP), which allows the regeneration of ADP from AMP (24). The combination of PAP and PPK therefore allowed complete regeneration of ATP from AMP (25). In an alternative approach, adenylate kinase (which catalyzes the reversible phosphorylation of AMP by ATP) was used in place of PPK. Coupling of this enzyme with PAP pushes the equilibrium towards ATP regeneration (Figure 1B) (26).

### The application of process modelling

Synthetic biology has long promised the use of well-characterized parts for the rational design of new metabolic pathways (27–29). However the use of modelling to optimize these *in vivo* reactions is yet to be fully realized (30). In contrast, modelling of enzymes *in vitro* can be fairly robust and offers solutions for reaction engineering (31). This could allow the combination of enzymes, for which validated mathematical models exist, to be rapidly engineered and tested *in silico* (2).

Indeed, in developing new biocatalytic processes the use of kinetic modelling is widely advocated, yet often not used in process development (32,33). The development of a kinetic model early on in the development process can be invaluable in cost/benefit or feasibility analysis (33). It permits evidence-based decision-making (34), and critically allows for the identification of bottlenecks and the quantification of process problems (e.g. feedback inhibition or inhibition by side-products) (6). Mechanistic models, which seek to describe enzyme mechanisms as accurately as possible, seek to understand a system as well as to predict it. These offer opportunities to develop substantial improvements or insights into the development process (31). A classic example is the use of Michaelis-Menten equations (35). Such integrated kinetic models of multi-enzyme processes are challenging because of the large number of kinetic parameters involved (34). Deterministic models for enzyme processes with two steps (plus cofactor regeneration) have been explored (34,36); alternative models have fitted kinetic parameters based on product concentrations (37,38). The most ambitious of these fitted parameters is for up to fifteen enzymes (39).

To demonstrate the use of CARs in multi-step cascade reactions, and investigate their use *in vitro*, we designed a reaction made up of an esterase, a CAR and an alcohol dehydrogenase (ADH) to hydrolyze and then reduce an ester to its corresponding alcohol. Methyl 4-toluate was chosen as a trial substrate. Whilst not directly industrially relevant, it acts as a good model for many industrially relevant compounds. The three step reaction models chemically simple reactions to supply an acid to the CAR, and to utilize the aldehyde product. Enzymes for cofactor regeneration and removal of an inhibitory by-product were added to provide an efficient reaction with sub-stoichiometric cofactor concentrations. We demonstrate that this results in an effective three step cascade for the ester to alcohol transformation incorporating seven enzymes over eight biochemical steps.

To highlight the modular nature of the enzyme toolbox for designing multistep reactions, we characterized each enzyme individually and established a mathematical model for the entire pathway. In constructing and testing the model for each step in the reaction we demonstrate the value of building a model in identifying process problems. Specifically, we were able to predict the need for PP_I_ removal, and a non-enzymatic reaction of an intermediate with a component of the reaction. We then performed optimization of the trial batch reaction, and achieved the target 90% yield with the minimal concentration of the enzymes. Our results demonstrate that a deterministic model can be robust for enzyme cascades with cofactor recycling involving at least seven enzymes. We further demonstrate that these models have great potential to quickly assemble novel enzyme catalyst networks (40).

## Results

### Expression and purification of enzymes

All of the enzymes used in this study were recombinantly prepared from *E. coli*, as fusions with a polyhistidine-tag. All of the proteins were purified by nickel affinity chromatography followed by size exclusion chromatography. Each enzyme was purified to >90% purity (Supplementary Figure 1 to Supplementary Figure 8).

### Designing a synthetic multi-step pathway

Enzymes with the correct substrate specificity to catalyze the esterase, CAR and ADH steps were identified from the literature. Where possible, thermostable enzymes were chosen to provide maximum operational stability. In the case of the CAR step, only moderate thermostability was possible due to the limited number of organisms that use these enzymes. An esterase from the hyperthermophile *Archaeoglobus fulgidus* (afEst2), a CAR from the moderate thermophile *Mycobacterium phlei* (mpCAR), and an ADH from the hyperthermophile *Aeropyrum pernix* (apADH) were chosen (Figure 1B) (41,42). The catalytic constants of these enzymes against the relevant substrates in the test pathway were determined (Supplementary Figure 19 to Supplementary Figure 38) or taken from relevant literature (references shown in Table 3).

To allow cofactor regeneration, a thermostable mutant of PTDH, capable of regenerating both NADH and NADPH, was chosen given the need for both of these cofactors in the pathway. For ADP regeneration from AMP, we identified a PAP enzyme from the thermophile *Thermodesulfobium narugense* (tnPAP) with 33 % identity to the previously characterized PAP from *Acinetobacter johnsonii* (43).

A thermostable PPT enzyme from *Thermosynechococcus elongatus* that had previously been characterized (44) was initially chosen to regenerate ATP from ADP. However, in our hands this enzyme gave very low activity. In its place, a thermostable AK enzyme from *Thermotoga neapolitana* (tnAK) was used, which had previously been characterized with high activity (45). We had previously identified that PP_I_ is a significant inhibitor of CARs (15). We therefore also determined the activity of a thermostable PPiase from *Thermus thermophilus* (ttPPiase; Supplementary Figure 33).

### Identifying an operational window

Thermostability and activity at different pHs were determined for each enzyme to define an operational window for the reaction (Supplementary Figure 39). Data for afEst2(42), mpCAR(15), apADH(46) and tnAK (45) were adapted from previous work. The effects of pH and temperature on the remaining enzyme activities were characterized (Supplementary Figures 11 to 14). A pH of 7.5 was chosen as a compromise for all enzymes, and ADH activity in the (undesired) oxidative direction was minimized at this pH. As we expected, the operational window for temperature is primarily limited by the mpCAR enzyme, as this has only moderate thermostability. A reaction temperature of 30 °C was chosen to ensure maximum activity of this enzyme.

### Reaction modelling

Mathematical models utilizing kinetics based on the steady state approximation were developed in isolation for each section of the multi-step reaction and validated before being combined into a multi-step process. These models were developed using Python scripts (Supplementary File 1), with equations appropriate for each enzyme. The rate equations are summarized in Table 1, which were used to construct differential equations for each reaction to be tested (Table 2). Each rate equation is based upon the steady-state approximation, and represents a unidirectional reaction (47). Where reactions are reversible, equations for both directions were included. The rationale for each equation used is detailed below. Kinetic parameters were either identified from the available literature or determined experimentally. An uncertainty analysis was carried out for each model, where possible, using bounds equal to the 95 % confidence intervals of each parameter (Table 3). In some cases where uncertainty was judged to be large due to incomplete knowledge, bounds of ± 50 % of the parameter were used. Starting concentrations were given bounds of ± 5 %, except in the case of polyphosphate where the length of the polyphosphate chain is unknown. In this case ± 25 % was used with the upper bound equal to the absolute concentration of phosphate units. Model predictions were tested by running small scale reactions in a thermomixer, with samples taken every 30 minutes and quenched with acetonitrile for analysis by HPLC.

**Table 1.**
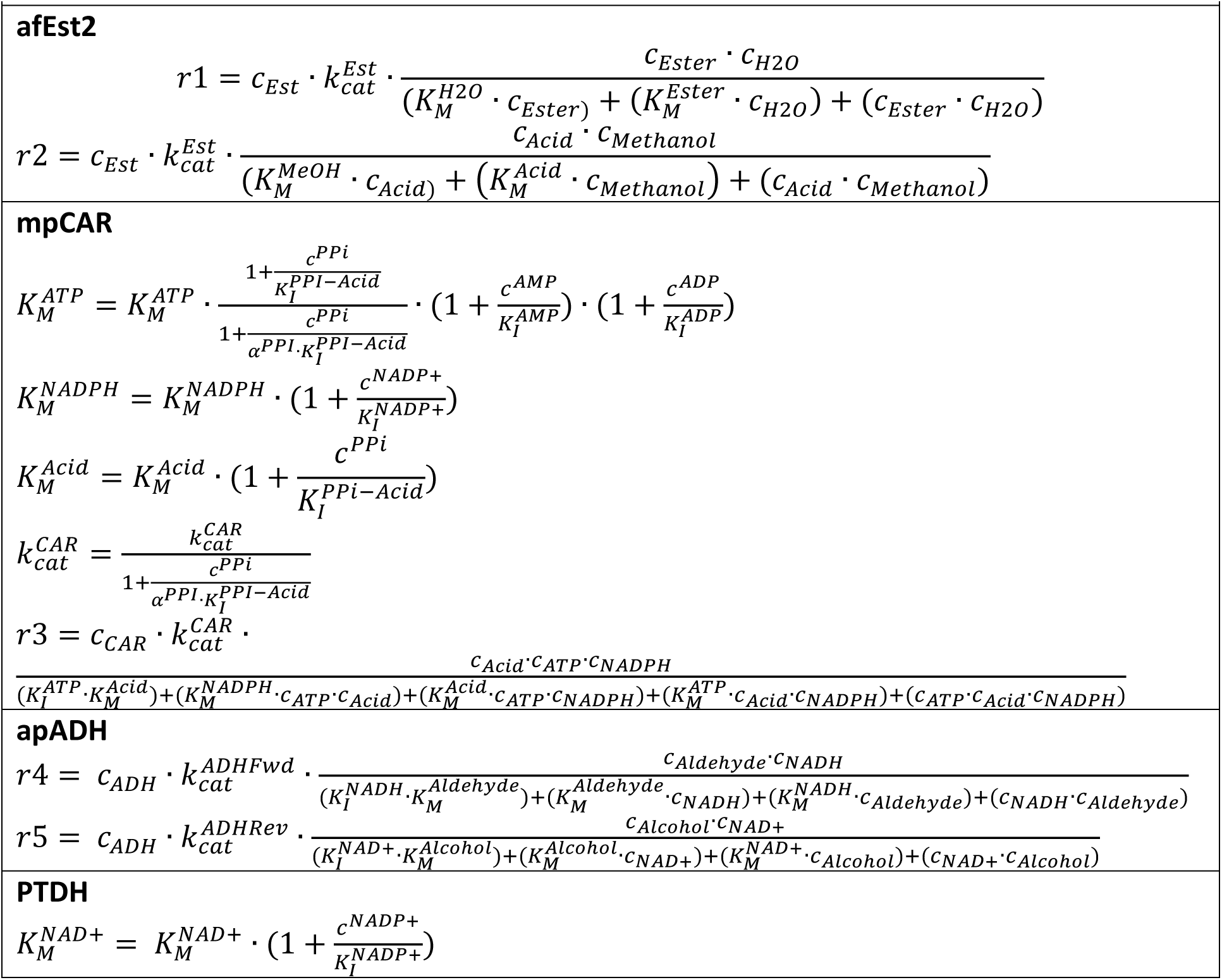
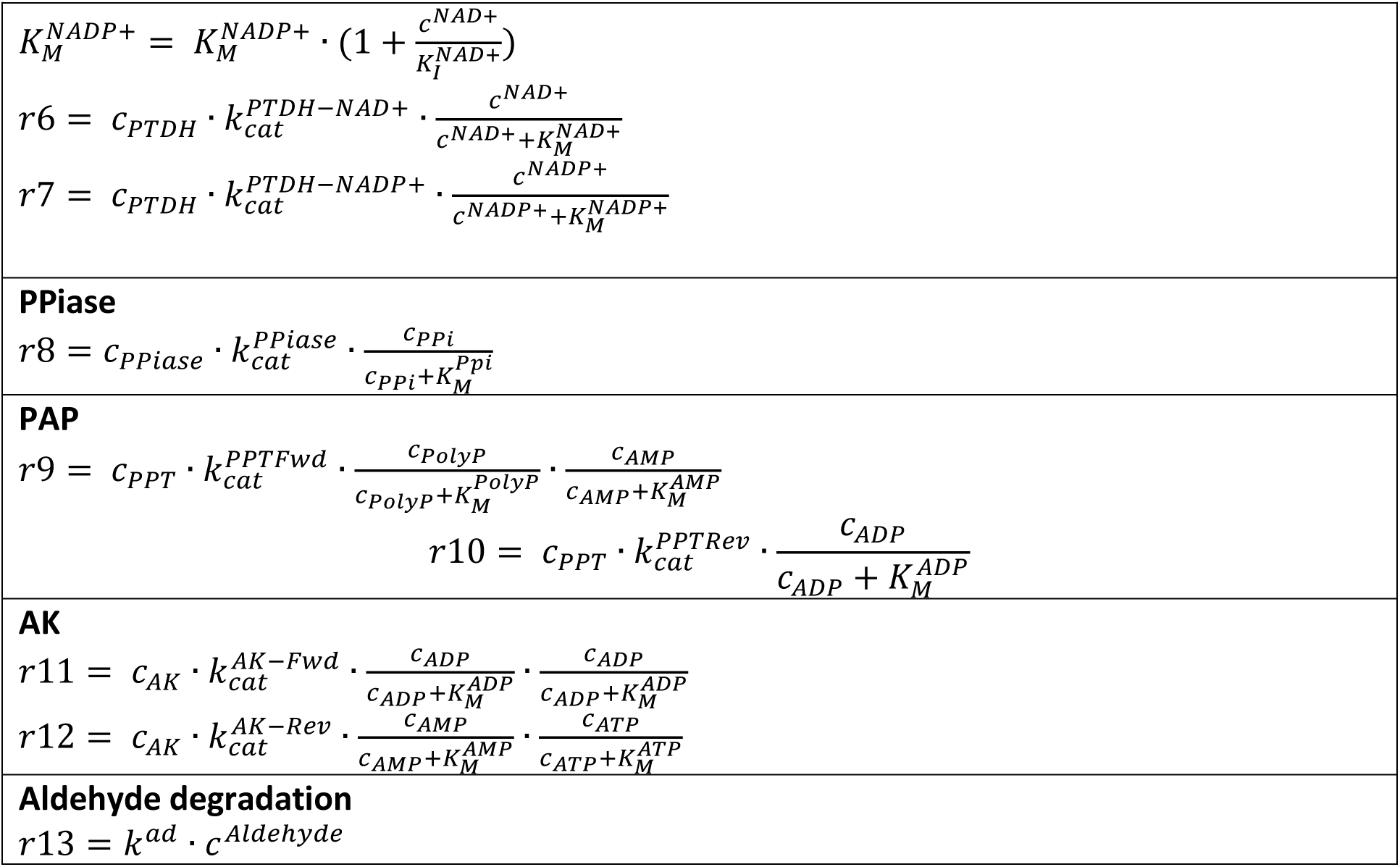
Kinetic equations. The rationale for the kinetic equations chosen for each enzyme and interacting partner are discussed in detail below. Aldehyde degradation was modelled as a first order process.

**Table 2.**
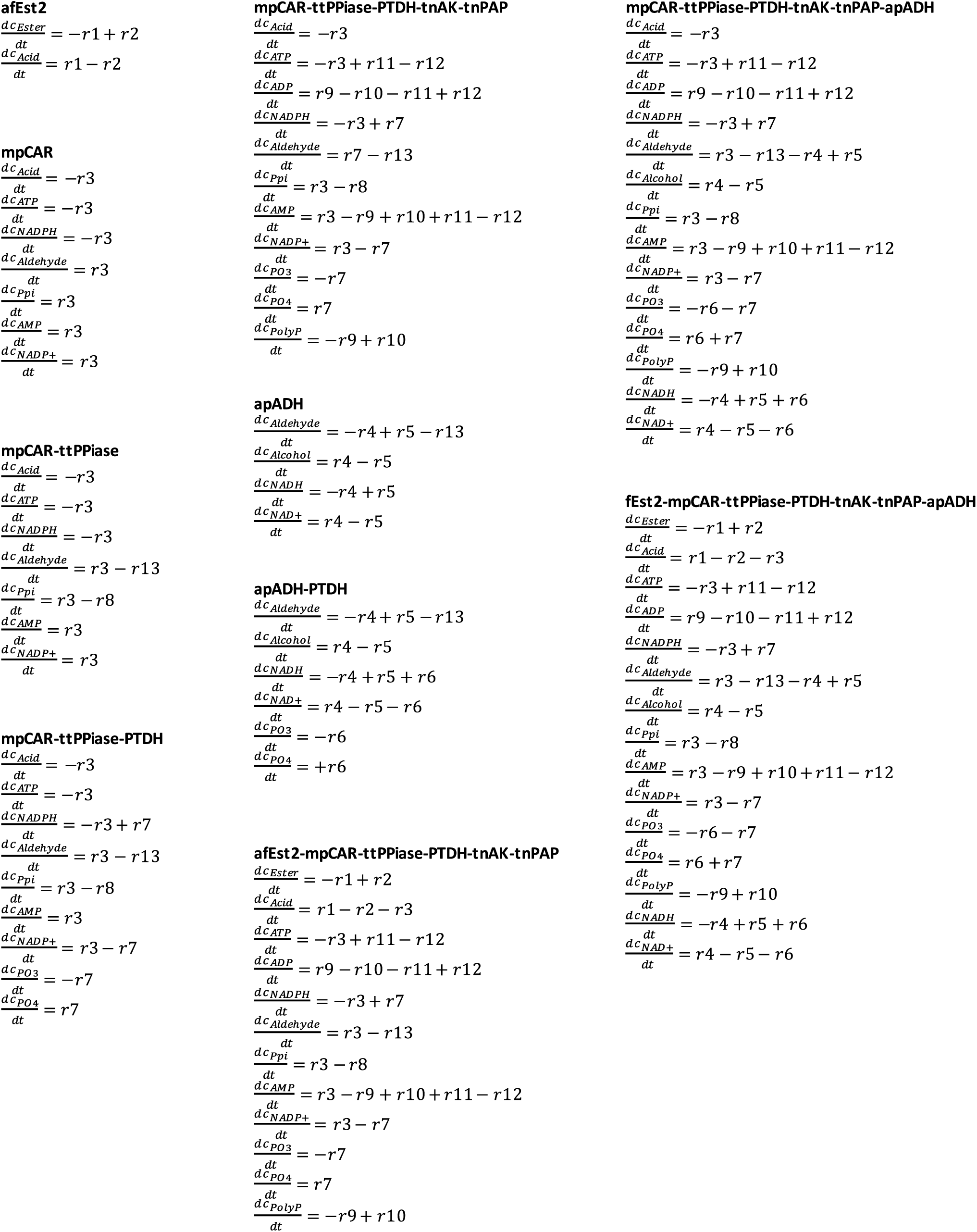
Differential Equations. Differential equations used to model changing substrate concentrations over time. *r*1 – *r*13 refer to the rate equations in Table 1.

**Table 3.** Kinetic parameters. Parameters were determined experimentally or obtained from the literature. Errors represent 95% confidence intervals where these could be experimentally determined or obtained from the literature. Where these could not be experimentally determined, errors of 50 % of the parameter value were conservatively assigned. In (a few) cases a reasonable estimate had to be made, marked with *. Where parameters were completely unknown, large uncertainty bounds were used, marked with **. *** For the esterase alone, and in the complete reaction respectively, reflecting data in Supplementary Figure 16.

| **afEst2 (42)** |  |
| --- | --- |
| $k_{cat-Fwd}$ (42) *** | $9.3 \pm 4.65$ or $12 \pm 6 \text{ min}^{-1}$ |
| $K_{M-Ester}$ (42) | $1,500 \pm 375 \mu\text{M}$ |
| $K_{M-H_2O}$ ** | $1,000 - 100,000 \mu\text{M}$ |
| $k_{cat-Rev}$ ** | $1 - 50 \text{ min}^{-1}$ |
| $K_{M-MeOH}$ ** | $1,000 - 100,000 \mu\text{M}$ |
| $K_{M-Acid}$ ** | $100 - 100,000 \mu\text{M}$ |
| **mpCAR (15)** |  |
| $k_{cat}$ | $200 \pm 20 \text{ min}^{-1}$ |
| $K_{M-Acid}$ | $1,500 \pm 320 \mu\text{M}$ |
| $K_{M-ATP}$ | $100 \pm 28 \mu\text{M}$ |
| $K_{I-ATP}$ | $40 \pm 34 \mu\text{M}$ |
| $K_{M-NADPH}$ | $30 \pm 8 \mu\text{M}$ |
| $K_{I-AMP}$ (15) | $10,000 \pm 1,800 \mu\text{M}$ |
| $K_{I-ADP}$ | $11,000 \pm 4,000 \mu\text{M}$ |
| $K_{I-NADP+}$ (15) | $143 \pm 16 \mu\text{M}$ |
| $K_{I-PPI-Acid}$ (15) | $340 \pm 80 \mu\text{M}$ |
| $K_{I-PPI-ATP}$ (15) | $220 \pm 100 \mu\text{M}$ |
| $\alpha\text{-PPI-ATP}$ (15) | $2.6 \pm 2.8$ |
| **apADH** |  |
| $k_{cat-Fwd}$ | $1.7 \pm 0.2 \text{ min}^{-1}$ |
| $k_{cat-Rev}$ * | $1.7 \pm 0.85 \text{ min}^{-1}$ |
| $K_{M-NADH}$ | $180 \pm 60 \mu\text{M}$ |
| $K_{M-NAD+}$ | $190 \pm 40 \mu\text{M}$ |
| $K_{I-NAD+}$ * | $185 \pm 92.5 \mu\text{M}$ |
| $K_{I-NADH}$ * | $185 \pm 92.5 \mu\text{M}$ |
| $K_{M-Aldehyde}$ | $350 \pm 120 \mu\text{M}$ |
| $K_{M-Alcohol}$ | $10,000 \pm 5,000 \mu\text{M}$ |
| **ttPpiase (72)** |  |
| $k_{cat}$ | $4,400 \pm 2,200 \text{ min}^{-1}$ |
| $K_{M-PPI}$ (72) | $500 \pm 250 \mu\text{M}$ |
| **PTDH** |  |
| $k_{cat-NAD+}$ | $637 \pm 16 \text{ min}^{-1}$ |
| $k_{cat-NADP+}$ | $342 \pm 16 \text{ min}^{-1}$ |
| $K_{M-NAD+}$ | $85 \pm 5 \mu\text{M}$ |
| $K_{M-NADP+}$ | $220 \pm 40 \mu\text{M}$ |
| **tnPAP** |  |
| $k_{catFwd}$ | $250 \pm 125 \text{ min}^{-1}$ |
| $K_{M-AMP}$ | $280 \pm 240 \mu\text{M}$ |
| $K_{M-PolyP}$ | $4,000 \pm 2,000 \mu\text{M}$ |
| $k_{catRev} (78)$ | $3.4 \pm 1.7 \text{ min}^{-1}$ |
| $K_{M-ADP} (78)$ | $8,300 \pm 4,150 \mu\text{M}$ |
| **tnAK (45,79)** |  |
| $k_{cat-ADP} (45)$ | $2,340 \pm 1,170 \text{ min}^{-1}$ |
| $k_{cat-AMP-ATP} (45)$ | $3,950 \pm 1,975 \text{ min}^{-1}$ |
| $K_{M-ADP} (79)$ | $91 \pm 45.5 \mu\text{M}$ |
| $K_{M-AMP} (79)$ | $38 \pm 19 \mu\text{M}$ |
| $K_{M-ATP} (79)$ | $51 \pm 25.5 \mu\text{M}$ |
| **Aldehyde side reaction** |  |
| $k$ | $0.00279 \pm 0.001395 \text{ min}^{-1}$ |

### Esterase reaction

The afEst2 has been recently characterized and kinetic parameters for the hydrolysis of methyl 4-toluate at 30 °C reported (42) (Table 3). As this characterization was carried out at pH 8.2, a medium level of ± 25 % uncertainty was associated with these parameters. Hydrolysis reactions are typically modelled using the irreversible, one substrate Michaelis-Menten equation. This model was tested against a small scale batch reaction (Figure 2A), which it was able to make a reasonable prediction. A reduced *χ*^2^ statistic was used to assess the model (48). A model fitted directly to the experimental data would be expected to achieve a *χ*^2^ of one; we interpreted models as strong where *χ*^2^ was less than fifteen, and less than a quadratic or simple linear model fitted to the experimental data. Here, *χ*^2^=2.9 (c.f. 6.3 and 32 for the alternative models), and so was interpreted as a good fit.

**Figure 2.**
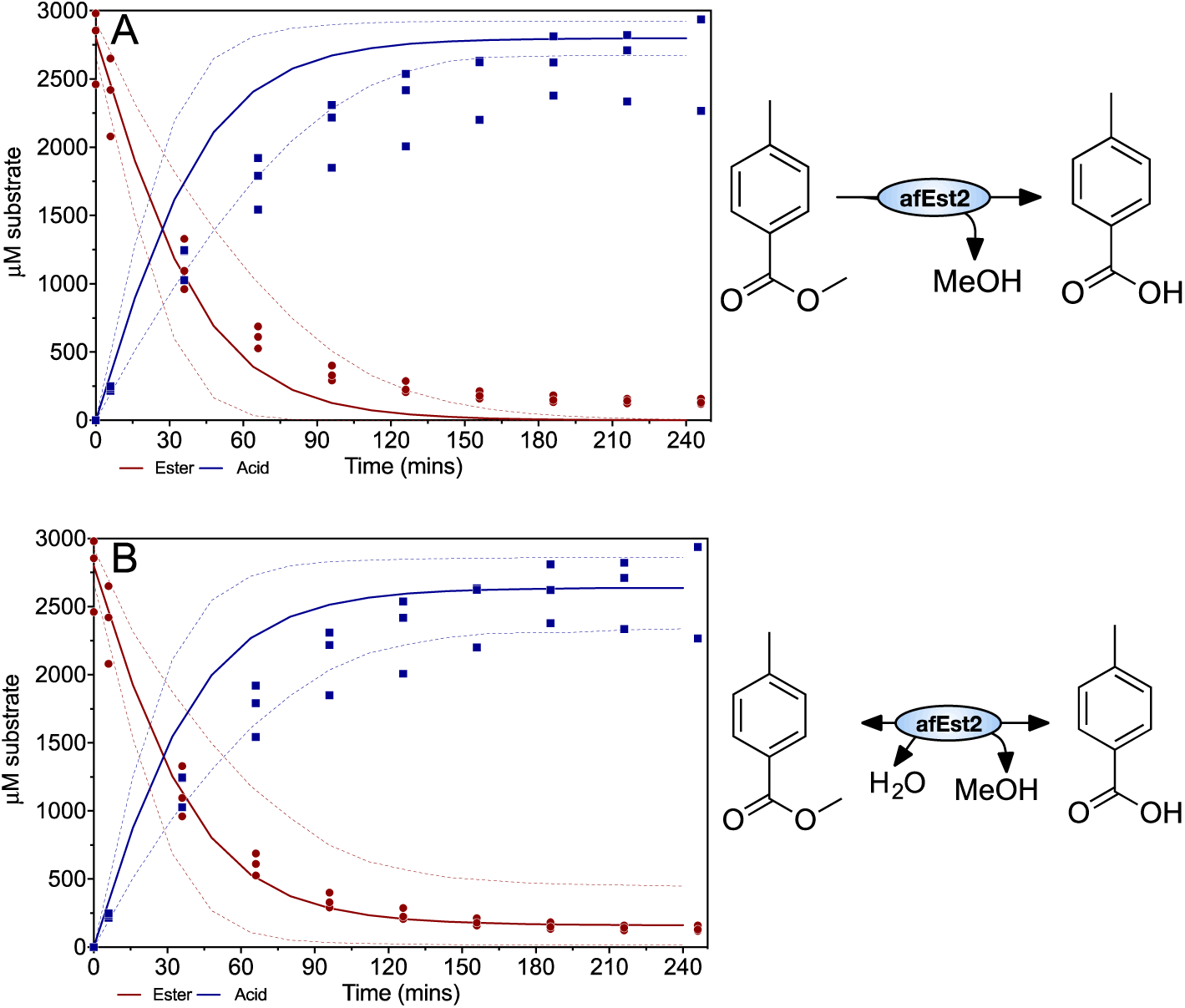
Validation of the esterase model. A reaction containing 10 μM afEst2 and 2,800 μM methyl 4-toluate (ester) was used to validate the model for afEst2. Ester and acid concentrations, measured every 30 minutes by HPLC, are shown as red circles and blue squares respectively. The model prediction is shown as the solid line in the same colors. *χ*^2^ = 2.9, 6.3, 32 for the deterministic, quadratic and linear models respectively; when the reverse reaction is included in the deterministic model, *χ*^2^ reduces to 0.82. Dashed lines represent the 5^th^ and 95^th^ percentile of the uncertainty analysis. Data show three experimental replicates for each point.

A. Esterase reaction modelled as an irreversible, one substrate reaction
B. Esterase reaction modelled as a reversible bi-bi reaction, with estimated parameters for the reverse direction.

However, on examination of the data, it is noticeable that the reaction proceeds faster than modelled, and that the reaction did not appear to go to completion. The characterization of this enzyme was performed in 2.5 mM buffer (necessarily as the assay detects a change in pH) (42), whilst all of our reactions used 100 mM buffer. We tested the effect of changing the buffer concentration on afEst2, and found that the enzyme is 55 % faster in 100 mM buffer (Supplementary Figure 15). To reflect this the *k*_*cat-Fwd*_ was increased by 55 % to 9.3 min^−1^, with ± 50 % uncertainty. While an irreversible reaction was initially assumed due to an excess of water, we calculated the *K*_*eq*_ for this reaction as 22, consistent with a noticeable reverse reaction. We therefore incorporated the reverse direction into our model (Table 1), with estimated parameters with large uncertainty bounds (Table 3) (49–51). The updated model predicted the batch reaction well, albeit with a larger degree of uncertainty (Figure 2B). A sensitivity analysis was carried out to look at the sources of uncertainty in this new model (Supplementary Figure 16). The revised model showed *χ*^2^=0.81, highlighting the improvement of the model.

### CAR reaction

The CAR enzymes have three substrates: ATP, NADPH and a carboxylic acid. The reaction proceeds in an ordered fashion, with the ATP activating the acid and an enzyme-acid conjugate being formed before NADPH binds (52). This reaction was therefore modelled using a three substrate rate equation, analogous to that of an aminoacyl tRNA synthetase in which a ternary complex must form before the third substrate can bind (53). Parameters were determined experimentally (Supplementary Figures 21 and 22).

A reaction to test the CAR model was set up, predicting a fast turnover of the acid into its derivative aldehyde (shown in grey, Figure 3A). Instead, after an initial burst, activity slowed considerably and the reaction was not complete even after four hours (*χ*^2^ = 200). We investigated the possibility of product inhibition and found that all of the products of the CAR reaction (PP_I_, AMP and NADP+) act as inhibitors (discussed in more detail in (15)). While AMP and NADP^+^ both act as competitive inhibitors which may be overcome by increasing the substrate concentration, PP_I_ acts as a mixed model inhibitor with respect to ATP, causing a large decrease in the apparent *k*_*cat*_. When the inhibition by these products was taken into account the model fitted the data quite well (Figure 3A; *χ*^2^ = 3.1). We noted that modelling the inhibition resulted in a large level of uncertainty.

**Figure 3.**
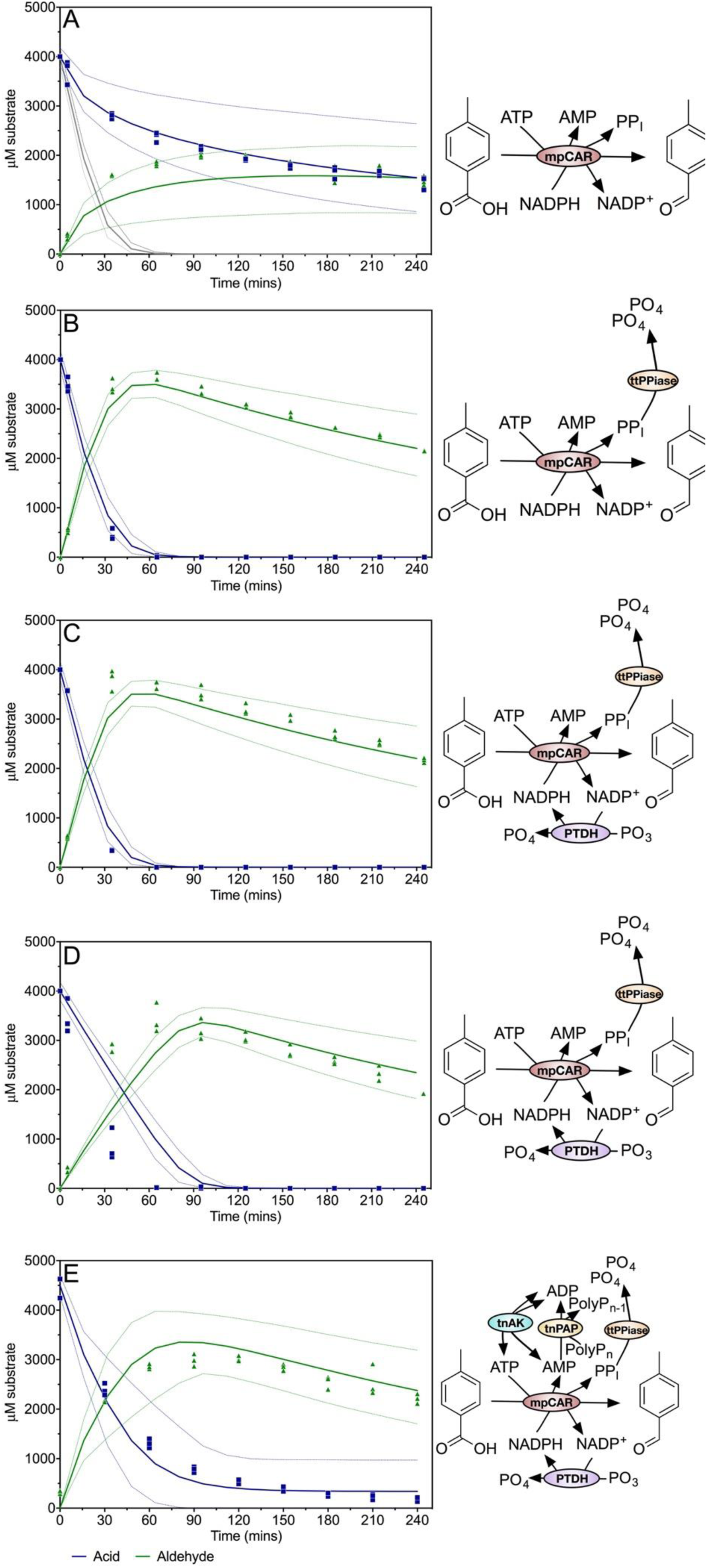
Validation of mpCAR and related enzyme models. Reactions were performed to validate the model for mpCAR, and the addition of cofactor regenerating and PP_i_ removing enzymes. Acid and aldehyde concentrations measured every 30 minutes by HPLC are shown as blue squares and green upwards facing triangles respectively. The model prediction is shown as the solid line in the same colors. Dashed lines represent the 5^th^ and 95^th^ percentile of the uncertainty analysis. Data show three experimental replicates for each point.

A. mpCAR alone. Reactions were initiated with 1 μM mpCAR, 8,000 μM ATP, 5,000 μM NADPH, 20,000 μM MgCl_2_, 4,000 μM 4-toluic acid. The initial model prediction, which did not take in to account any product inhibition, is shown in grey.
B. mpCAR-ttPpiase. Reactions were initiated with 1 μM mpCAR, 1 μM ttPpiase, 8,000 μM ATP, 5,000 μM NADPH, 20,000 μM MgCl_2_, 4,000 μM 4-toluic acid.
C. mpCAR-ttPpiase-PTDH. Reactions were initiated with 1 μM mpCAR, 1 μM ttPpiase, 1 μM PTDH, 8,000 μM ATP, 500 μM NADPH, 20,000 μM MgCl_2_, 20,000 μM PO_3_, 4,000 μM 4-toluic acid.
D. mpCAR-ttPpiase-PTDH (low NADPH). Reactions were initiated with 1 μM mpCAR, 1 μM ttPpiase, 1 μM PTDH, 8,000 μM ATP, 50 μM NADPH, 20,000 μM MgCl_2_, 20,000 μM PO_3_, 4,000 μM 4-toluic acid.
E. mpCAR-ttPpiase-PTDH-tnPAP-tnAK. Reactions were initiated with 0.4 μM mpCAR, 1 μM ttPpiase, 1 μM PTDH, 3 μM tnPAP, 1 μM tnAK, 1,250 μM ATP, 500 μM NADPH, 20,000 μM MgCl_2_, 20,000 μM PO_3_, 6,000 μM polyphosphate, 4,000 μM 4-toluic acid. Observed reduced *χ*^2^ values (for deterministic, fitted quadratic and fitted linear models): A: 3.1, 2.3, 21; B: 2.4, 220, 780; C: 160, 1300, 1400; D: 1200, 10, 1100; E: 11, 30, 210.

### ttPPiase removes CAR inhibition by PP_I_

In order to alleviate inhibition by PP_I_ on the CAR enzyme, an inorganic pyrophosphatase from *T. thermophilus* (ttPPiase) was added to the reaction. This enzyme shows exceptional thermostability and retained over 90 % activity after a 30 minute incubation at 95 °C (Supplementary Figure 39). Pyrophosphatase activity was measured at a saturating concentration of PP_I_ (5 mM) and the rate taken to be the *k*_*cat*_. The enzyme was modelled using the one substrate Michaelis-Menten equation, and using the *K*_*M*_ determined in a recent study on this enzyme (54). The addition of the ttPPiase alleviated the majority of the inhibitory effects seen previously and the CAR reaction went to completion within one hour (Figure 3B).

### Aldehyde side reaction

Once the CAR reaction was complete, aldehyde concentration decreased over time (Figure 3B). Aldehydes can react with free amines; and benzaldehyde in particular has a documented specific reaction with Tris (Supplementary Figure 17) (55,56). We verified that 4-methylbenzaldehyde reacts in a similar manner with Tris, and validated the product by mass spectrometry (Supplementary Figure 18). As we validated this side reaction after completing all other experiments, we accounted for this reaction using a one phase decay equation. This was fitted to the decrease in aldehyde concentration, and an equation for the rate of aldehyde side reaction constructed (Table 1 and Table 3). This resulted in a very good fit of the aldehyde concentration to the data (*χ*^2^ = 2.4).

### NAD(P)H regeneration using PTDH

The regeneration of NADPH using PTDH was then added to the CAR step. Although PTDH shows sequential ordered kinetics with NAD^+^ binding before phosphite, the effect of phosphite concentration was not modelled as all experiments were performed at saturating phosphite concentrations. The one substrate Michaelis-Menten equation was therefore used for reactions with either NAD^+^ or NADP^+^ as the substrate. NAD^+^ and NADP^+^ act as competitive inhibitors of each other with *K*_*I*_ values equal to their respective *K*_*M*_ values, and this inhibition was added to the model (Table 1). Kinetic parameters for NAD^+^, NADP^+^ were determined at assay conditions by following the generation of NADH or NADPH at 340 nm (Table 3; Supplementary Figures 31 and 32).

The NADPH regeneration was tested by attempting reduction of 4000 μM 4-toluate by CAR with only 500 μM or 50 μM NADPH, together with the PTDH regeneration system. Both reactions went to completion with far less than stoichiometric NADPH. The reaction containing 500 μM NADPH fitted the model predictions reasonably well (Figure 3C; *χ*^2^ = 160, in this case driven by a single anomalous data point). When only 50 μM NADPH was used, a much slower reaction was predicted by the model. However, only a small difference was observed experimentally compared to the use of 500 μM NADPH (Figure 3D; *χ*^2^ = 1,200). We considered that some form of substrate channeling (57,58) may be taking place between the CAR and PTDH resulting in a lower apparent *K*_*M*_ for NADP^+^ by PTDH.

### ATP regeneration using tnPAP and tnAK

tnPAP showed good PAP activity, and kinetic parameters for polyphosphate and AMP were determined experimentally (Supplementary Figures 35 to 37), and a bi-substrate equation constructed. An adenylate kinase from *Thermotoga neapolitana* has been reported to possess excellent thermostability, with kinetic parameters similar to those of the characterized *E. coli* adenylate kinase. These parameters were used to model the adenylate kinase. However, testing of this enzyme in a CAR coupled reaction revealed a significantly lower turnover number (Supplementary Figure 40). Furthermore, the magnesium concentration has been shown to be instrumental in controlling the equilibrium catalyzed by AK. Since the free magnesium concentration is not known due to chelation by polyphosphate and other compounds, this effect was not taken into account. For this reason, the AK reaction is difficult to model accurately in the context of the multistep pathway. Therefore, a simple bi-substrate equation was used to describe it. Large uncertainty bounds were used for all the parameters in the AK reaction as a consequence. With all the cofactor regeneration systems in place, the reaction proceeded almost to completion with far less than stoichiometric cofactors, and fitted well to the model (Figure 3E; *χ*^2^ = 11).

### ADH reaction

The kinetic parameters for both the forward and reverse directions of the apADH catalyzed reactions were determined experimentally. The *K*_*M*_ for 4-tolyl alcohol (for the reverse oxidative reaction) could not be accurately determined, as the reaction is not saturated at the highest concentrations that the substrate is soluble at (Supplementary Figures 29 and 30). An estimated *K*_*M*_ of 100 mM was used (Table 3). Consequently, *k*_*cat-Rev*_ was also difficult to determine and a value similar to *k*_*cat-Fwd*_ was assumed. A two substrate steady state sequential rate equation was used to describe both the forward and reverse reactions separately, using an estimated parameter for the *K*_*I*_ for NAD+ and NADH based on their determined *K*_*M*_ values.

Reactions featuring only apADH as well as with PTDH for NADH regeneration (using far less than stoichiometric NADH) were used to validate the model, which predicted the rate of reduction well in both cases (Figure 4; *χ*^2^ = 7.0). A sensitivity analysis of the final alcohol concentration in both these reactions showed the *K*_*I*_ for NAD+ and NADH, and *k*_*cat-Rev*_ to have little effect (Supplementary Figure 42).

**Figure 4.**
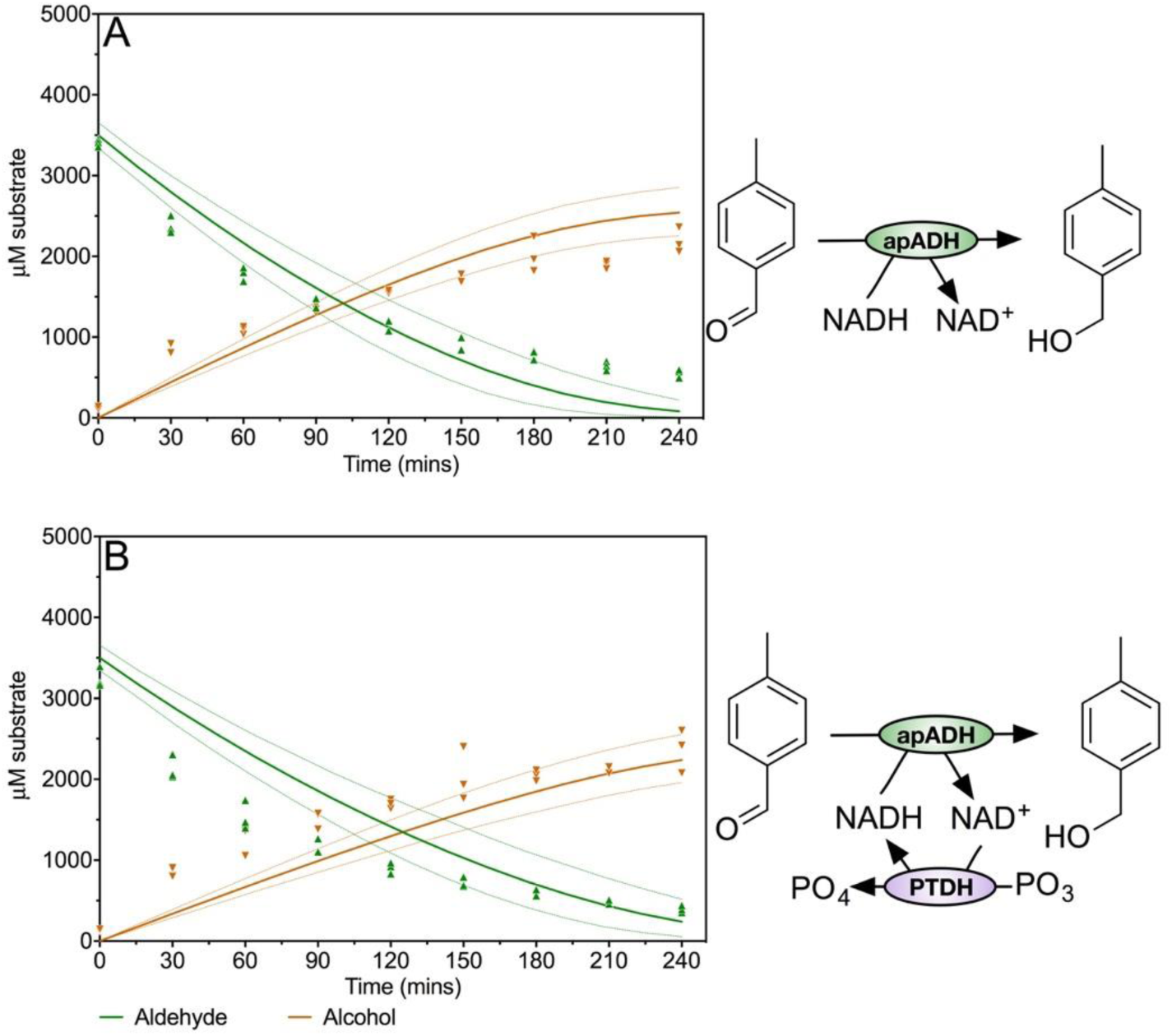
Validation of ADH and ADH-PTDH models. Reactions used to validate the model for apADH, and cofactor regeneration by PTDH. Aldehyde and alcohol concentrations measured every 30 minutes are shown as green upwards facing triangles and orange downward facing triangles respectively. The model prediction is shown as the solid line in the same colors. Dashed lines represent the 5^th^ and 95^th^ percentile of the uncertainty analysis. Data show three experimental replicates for each point.

A. apADH only. Reactions were initiated with 10 μM apADH, 5,000 μM NADH, 3,500 μM 4- tolualdehyde.
B. apADH-PTDH. Reactions were initiated with 10 μM apADH, 1 μM PTDH, 20,000 μM PO_3_, 500 μM NADH, 3,500 μM 4-tolualdehyde. Observed reduced *χ*^2^ values (for deterministic, fitted quadratic and fitted linear models): A: 13, 7.9, 45; B: 7.0, 3.9, 35.

### Building a multi-step enzyme cascade

Once the models for each of the esterase, CAR and ADH steps were independently validated with cofactor regeneration, multi-step reactions were constructed. The CAR step was tested in combination with first the esterase, and then ADH (Figure 5A, B; *χ*^2^ = 10, 4.8 respectively).

**Figure 5.**
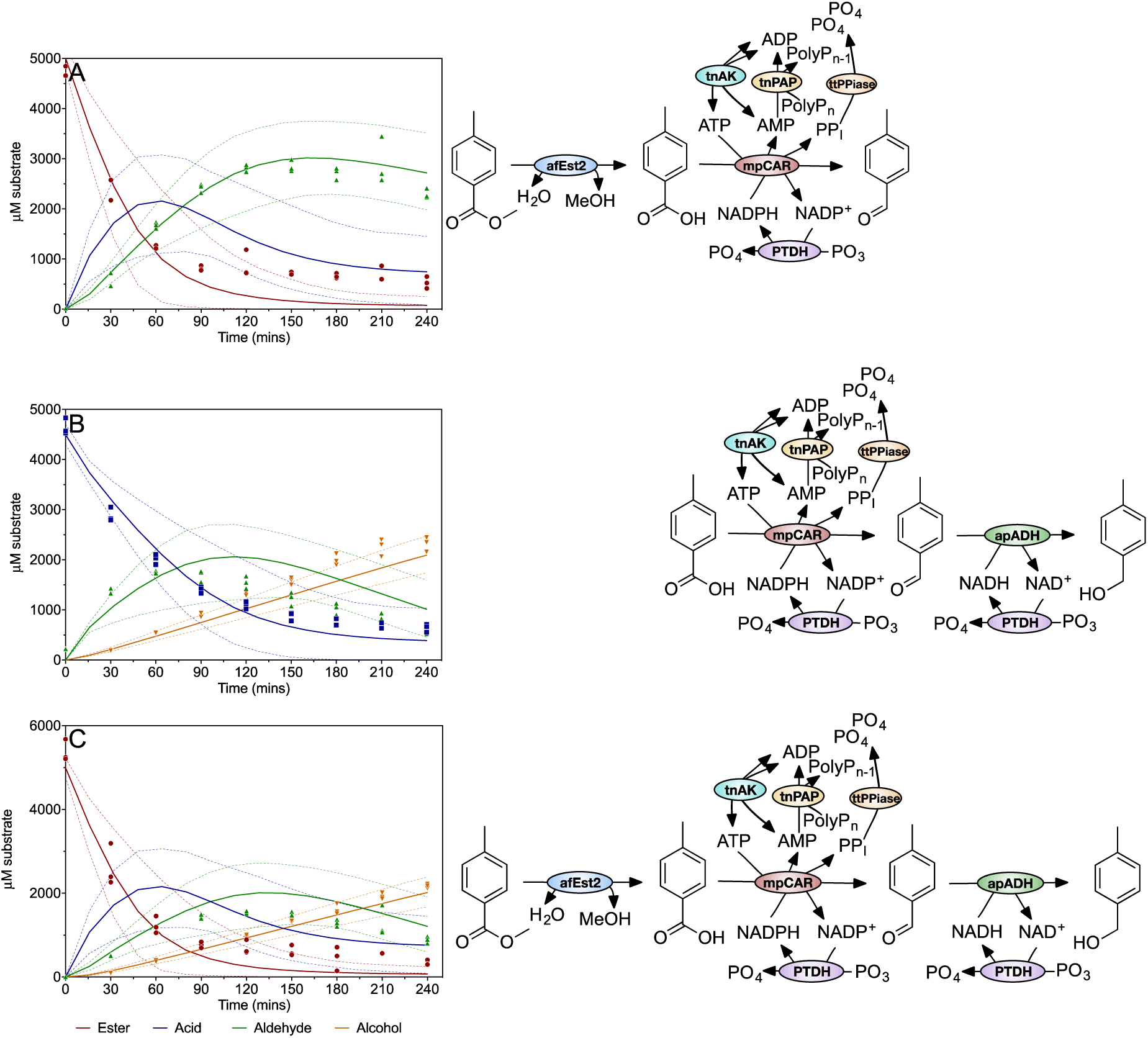
Validations of combinations of the esterase, CAR and ADH reactions. Reactions combining the CAR step with the esterase, ADH or both were constructed to test the model for these multi-step reactions. Ester, acid, aldehyde and alcohol concentrations measured every 30 minutes are shown as red circles, blue squares, green upwards facing triangles and orange downward facing triangles respectively. The model prediction is shown as the solid line in the same colors. Dashed lines represent the 5^th^ and 95^th^ percentile of the uncertainty analysis. Data show three experimental replicates for each point.

A. afEst2-mpCAR-ttPpiase-PTDH-tnPAP-tnAK. Reactions were initiated with 10 μM afEst2, 0.4 μM mpCAR, 1 μM ttPPiase, 1 μM PTDH, 3 μM tnPAP, 1 μM tnAK, 5,000 μM methyl 4-toluate, 500 μM NADPH, 1,250 μM ATP, 20,000 μM phosphite, 6,000 μM polyphosphate, 20,000 MgCl_2_.
B. mpCAR-ttPpiase-PTDH-tnPAP-tnAK-apADH. Reactions were initiated with 0.4 μM mpCAR, 1 μM ttPPiase, 1 μM PTDH, 3 μM tnPAP, 1 μM tnAK, 10 μM apADH, 4,500 μM 4-toluic acid, 500 μM NADPH, 500 μM NADH, 1,250 μM ATP, 20,000 μM phosphite, 6,000 μM polyphosphate, 20,000 MgCl_2_.
C. afEst2-mpCAR-ttPpiase-PTDH-tnPAP-tnAK-apADH. Reactions were initiated with 10 μM afEst2, 0.4 μM mpCAR, 1 μM ttPPiase, 1 μM PTDH, 3 μM tnPAP, 1 μM tnAK, 10 μM apADH, 5,000 μM methyl 4-toluate, 500 μM NADPH, 500 μM NADH, 1,250 μM ATP, 20,000 μM phosphite, 6,000 μM polyphosphate, 20,000 MgCl_2_. Observed reduced *χ*^2^ values (for deterministic, fitted quadratic and fitted linear models): A: 7.8, 150, 5200; B: 4.8, 8.9, 47; C: 5.4, 7.5, 24.

In each case the model performed well in predicting the productivity of the reaction. However, the hydrolysis reaction catalyzed by afEst2 proceeded faster than was expected (Figure 5A and 5C). We considered that one or more reagents for other reactions might increase the afEst2 activity. Addition of all of the final reaction reactants caused a 30% increase in rate, compared to the rate in 100 mM Tris (Supplementary Figure 15). Further experiments showed that no individual component accounted for this, implying a salting-in effect (Supplementary Figure 15). To accommodate this increase and the uncertainty surrounding it in to the modelling, we increased the esterase *k*_*cat*_ to two times the parameter value for the combined reactions, and increased the uncertainty to 50% of this value. Higher levels of uncertainty in the afEst2 step predictions are therefore observed (Figure 5A and 5C).

### Optimization of enzyme concentrations by a genetic algorithm

For most industrial processes, the cost of components is one critical factor in determining the economic feasibility of an approach. We therefore aimed to demonstrate that this multistep cascade could be optimized. We aimed to increase the overall yield of alcohol, whilst minimizing the overall concentration of enzymes used. We custom built a genetic algorithm (Supplementary Figure 43) to perform this optimization. Our algorithm generates random solutions for the concentration of each enzyme, and scores the quality of the solution against targets of 90% yield of alcohol, with the lowest total enzyme concentration. The highest scoring solutions are retained, and used to generate a new population of solutions. 25 cycles of this process were sufficient for the population to achieve a stable solution. Enzyme concentrations of 9.65 μM afEst2, 0.73 μM CAR, 24.75 μM ADH, 0.07 μM PPiase, 0.43 μM PTDH 0.15 μM AK and 0.57 μM PAP were identified as the lowest enzyme concentration pathway capable of achieving 90 % yield in four hours. This achieved a more than two-fold increase in yield, whilst reducing the concentration of six of the seven enzymes used.

To test the optimized pathway, reactions were performed at the reagent concentrations proposed by the optimization algorithm (Figure 6). The model fitted the data well (*χ*^2^ = 9.5). The final yield of alcohol was slightly lower than expected reaching a final yield of only 3,000 μM as opposed to the 3,500 μM predicted by the model. However, the optimized pathway performed significantly better in productivity compared to the non-optimized pathway.

**Figure 6.**
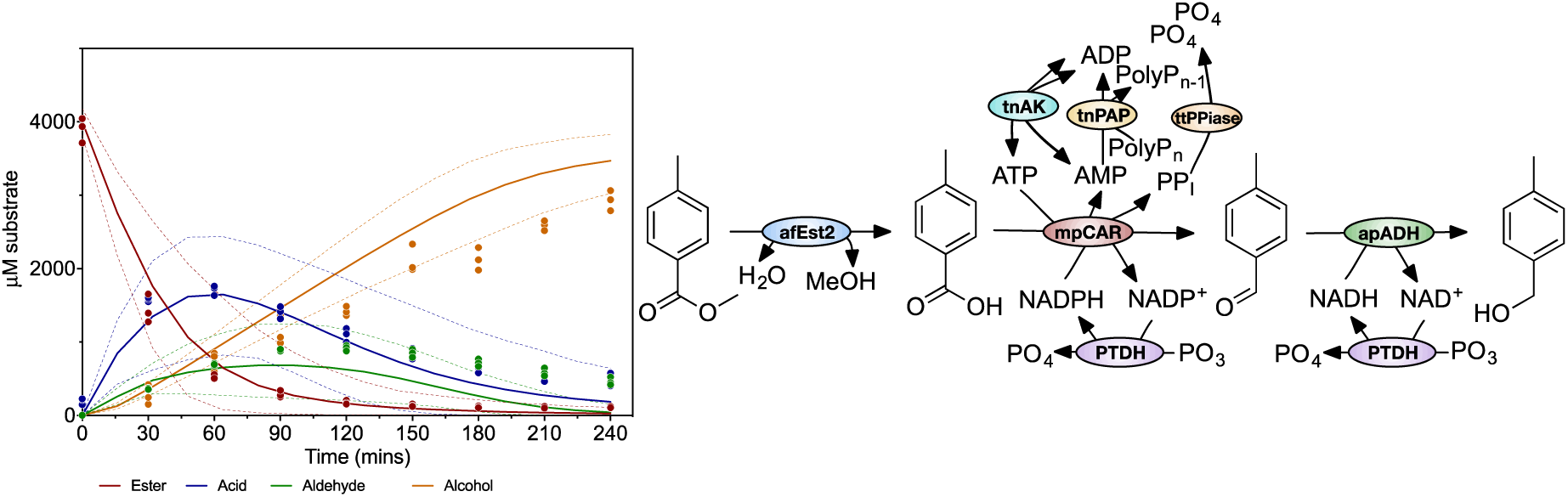
Validations of the optimized complete reaction. A reaction using enzyme concentrations predicted by a genetic algorithm to give the lowest cost reaction, whilst hitting a target of 90 % alcohol yield, was performed. Ester, acid, aldehyde and alcohol concentrations measured every 30 minutes are shown as red circles, blue squares, green upwards facing triangles and orange downward facing triangles respectively. The model prediction is shown as the solid line in the same colors. Dashed lines represent the 5^th^ and 95^th^ percentile of the uncertainty analysis. Data show three biological replicates for each point. The reactions were initiated with 9.65 μM afEst2, 0.73 μM mpCAR, 0.07 μM ttPPiase, 0.43 μM PTDH, 0.57 μM tnPAP, 0.15 μM tnAK, 24.7 μM apADH, 4,000 μM methyl 4-toluate, 500 μM NADPH, 500 μM NADH, 1,250 μM ATP, 20,000 μM phosphite, 6,000 μM polyphosphate, 20,000 μM MgCl_2_. Observed reduced *χ*2 values (for original deterministic, modified deterministic, fitted quadratic and fitted linear models): 9,5, 31, 79.

### Sensitivity analysis comparison of the optimized vs non-optimized pathways

As well as delivering optimal performance at lowest cost, it is important that industrial processes are robust, and not easily affected by issues with a single reagent. Therefore, a Sobol sensitivity analysis was carried out on the model for the optimized reaction to determine the main sources of the large uncertainty in the final alcohol concentration (59) (Figure 7). This revealed that the final alcohol yield was most sensitive to the parameters for tnAK in this optimized reaction (>30% of total uncertainty associated with these). Other parameters with a substantial contribution included polyphosphate concentration, CAR inhibition by PP_I_, and the rate of aldehyde degradation. These parameters were very different from those observed in the non-optimized reaction (Figure 7). This indicated that, whilst the optimization had been highly successful in delivering greater productivity at lower cost, this came at the cost of increased sensitivity to some parameters related to individual enzymes (resulting in a 3.5 fold increase in the 95% CIs of the final yield).

**Figure 7.**
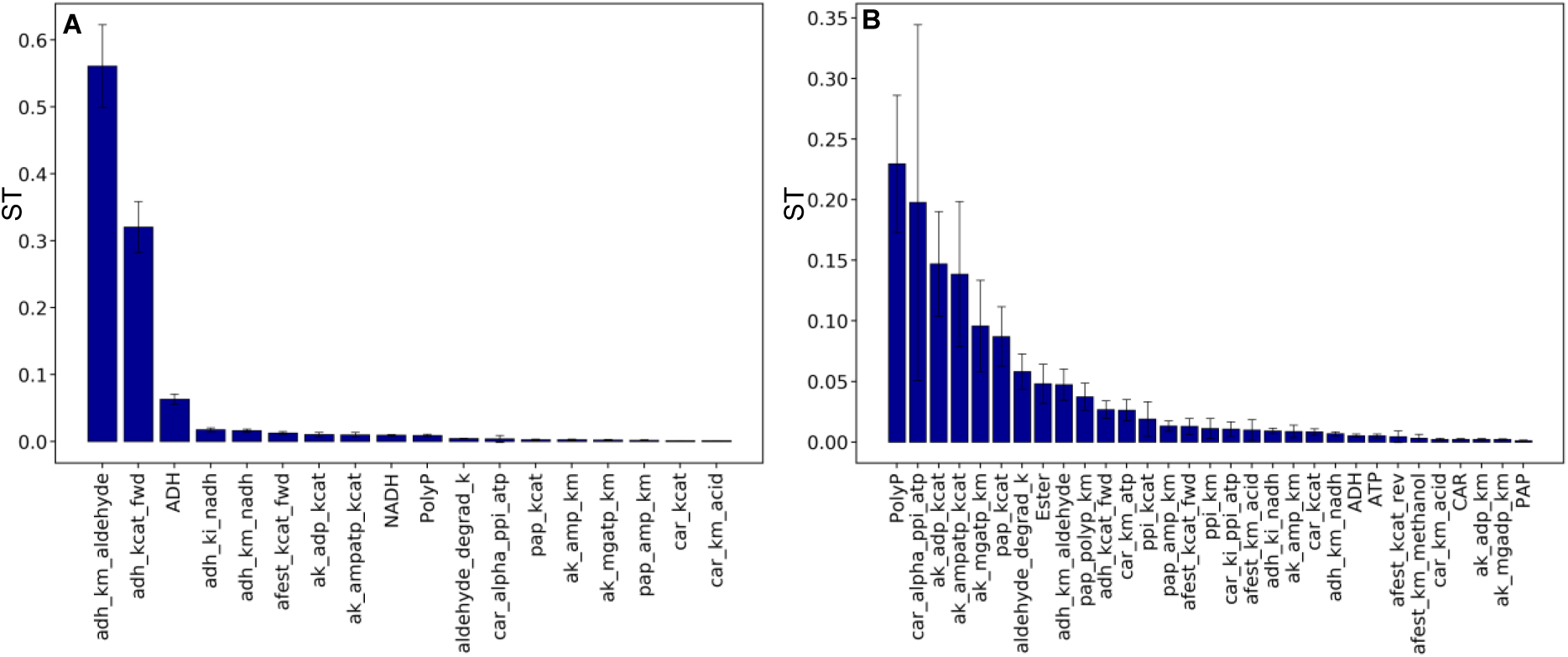
Sensitivity analysis of the modelled pre- and post-optimization batch reaction. The total sensitivity indices (ST) are shown which take into account 1^st^ order and all other interactions. Total of all STs for a reaction sum to 1. Sensitivity is in reference to the uncertainty in the final 4-tolyl alcohol concentration. Error bars show the 95 % confidence intervals. The sum of all sensitivity indices’ should equal 1. Parameters with a ST of less than 0.001 are not shown. **A**: Sensitivity analysis of the pre-optimization complete reaction, shown in figure 5C. **B**: Sensitivity analysis of the optimized complete reaction, shown in figure 6.

## Discussion

### Multi-step *in vitro* biocatalysis

Among several other advantages, the use of isolated enzymes allows reactions to be easily controlled and subsequently optimized quickly and easily. We have demonstrated the use of CARs for multi-step *in vitro* biocatalysis, making use of mechanistic modelling to understand the dynamics of multiple enzymes working in concert. The modular nature of constructing models for each enzyme allows a systems biology approach to be taken for the creation of new cascades, with the effects of enzyme module addition or removal being predictable *in silico*. Furthermore, establishing a mathematical model for this seven enzyme *in vitro* cascade facilitates its understanding. Key advantages include the ability to exploit the model for the evaluation of different process options, identification of bottlenecks and optimization of the reaction.

### Modelling drives hypothesis generation

Most importantly, modelling was critical in identifying the inhibition of CAR by PP_I_ (Figure 2). This has since been independently shown by Kunjapur et al. (19). This demonstrates one benefit of mechanistic modelling over a purely data driven approach. Upon observing the slower rate of the CAR reaction compared to the model expectations, we were able to focus efforts on determining the cause of this (15). We were then able to test the addition of the PPiase enzyme *in silico* before committing to the process of preparing this enzyme for inclusion in the cascade. It is possible that other enzymes might have unidentified inhibitors or activators in the cascade. However, our modeling showed no evidence of these, and so provides a data-led rationale for avoiding the costly testing of all possible pairwise interactions.

Testing our observations against model predictions revealed some other interesting phenomena. The inclusion of PTDH allowed the CAR reaction to proceed more effectively than the model predicted at low NAPDH concentrations (Figure 3D). More detailed modelling would be required to fully understand this. However, this observation might be a result of substrate shuttling or channeling between PTDH and the reductase domain of the CAR enzyme (60). It could be expected that the local concentration of NADP^+^ around the CAR enzyme might be higher, and that here PTDH would work more efficiently providing a larger local supply of NADPH (61). Possibly PTDH could even localize near the CAR for this reason.

### The use of CARs *in vitro*

CARs have previously been used in whole-cell biocatalysis where ATP and NADPH can be regenerated by the host metabolism (62). Using CARs as isolated enzymes allows the design of an efficient process, and the inclusion of cofactor regeneration makes this an economically competitive alternative to whole-cell biocatalysis.

PTDH accepts two substrates, NAD^+^ and NADP^+^, which compete for the enzyme’s active site. In our optimized reaction both NAD^+^ and NADP^+^ were set to 500 μM and only PTDH concentration was optimized. In this situation NAD^+^ is regenerated more effectively as PTDH has a lower *K*_*M*_ and higher *k*_*cat*_ for this substrate. Possibly the PTDH reaction could be optimized further by allowing the ratio of NAD^+^: NADP^+^ to be altered in the optimization procedure, to balance NADH and NADPH regeneration to the requirements of the reaction. The evidence for substrate shuttling in these reactions also offers the possibility to further reduce the concentrations of expensive cofactors.

### The ATP regeneration system is challenging to model

The reaction catalyzed by AK is critically regulated by magnesium (63). While AK has been shown to follow a random Bi Bi mechanism (64), knowledge of the free and bound magnesium concentrations is required to implement this mechanism as a rate equation. As this is impractical, we were forced to only approximate the AK reaction using a bi-substrate equation (Table 1), with high levels of uncertainty associated with each parameter. The uncertainty analysis performed easily captures the potential positions of equilibrium that would be modelled using a more complex but accurate rate equation, facilitating the use of the approximation (63). Future iterations of the model could seek to more accurately model the free and bound magnesium concentrations (65).

The reaction catalyzed by PAP is also difficult to model. Polyphosphate cannot be added at a saturating concentration due to chelation of magnesium, and the chain lengths and concentrations of the various polyphosphate species are difficult to determine. Since both polyphosphate chain length and concentration affect reaction kinetics, the reaction was again approximated using a bisubstrate equation with high levels of uncertainty associated with the polyphosphate concentration (Table 1).

### Reaction optimization

Optimization of the batch reaction, used to validate the model, demonstrates its potential for exploring new process options. Optimization of the reaction resulted in just enough afEst2 to hydrolyze all of the methyl 4-toluate in the time available, and just enough ApADH to achieve 90 % of the theoretical maximum of 4-tolyl alcohol at 4 hours (Figure 6). In contrast, 4-toluic acid and 4-tolualdehyde concentrations were maintained at a low but steady concentration allowing maximum productivity by all enzymes for the entire reaction. Enzyme concentrations were minimized and productivity maximized successfully.

In comparing the uncertainty in the pre-optimization (Figure 5C) and optimized (Figure 6) complete reactions, it is clear that uncertainty in the optimized reaction has increased significantly (Figure 7). Minimizing the concentration of all enzymes so that they are each close to being rate limiting has likely increased the uncertainty, as changes in a greater number of parameters can impact on the rates.

The pre-optimization reaction did show uncertainty in 4-toluic acid and 4-tolualdehyde concentrations, yet this does not translate to a high degree of uncertainty in the 4-tolyl alcohol concentration. When 4-tolualdehyde concentration is high enough above its *K*_*M*_ for ApADH, the main driver for 4-tolyl alcohol concentration uncertainty is the rate of the ApADH reaction (principally the adh_kcat_fwd parameter; Figure 7A). However when 4-tolualdehyde is maintained at the lower concentrations in the optimized reaction, this concentration has a greater impact on the rate of 4-tolyl alcohol production. Many of the parameters causing uncertainty in 4-tolualdehyde concentration are then responsible for the larger degree of uncertainty in 4-tolyl alcohol production (Figure 7B). This suggests that the sensitivity of product 4-tolyl alcohol production could be kept low by optimizing the 4-tolualdehyde concentrations to be maintained above the ApADH *K*_*M*_ for this intermediate. Future optimization could include an additional objective for the minimization of uncertainty in the final 4-tolyl alcohol concentration. This would allow costs to be minimized whilst also minimizing the uncertainty in the final yield. However the inclusion of two competing objective functions would make the optimization process more complex, likely requiring the generation and manual evaluation of a set of Pareto optimal solutions (66).

We estimate the cost of our optimized reaction (Figure 6), as approximately $0.0014 (for a 500 μL reaction; or $6 per gram of a 150 Da product; Supplementary Data), when performed at a scale to benefit from bulk prices (67). Further reductions in the cost would likely be achieved at high scales. Re-use of the enzymes (e.g. by enzyme immobilization) would significantly reduce enzyme costs; and cofactor costs could be minimized using the cheaper spent cofactors as the starting material. This would make the cascade competitive for the production of fine chemicals (e.g. perfumes or pharmaceuticals).

### Uncertainty and sensitivity analysis facilitate the use of approximations

The use of uncertainty analysis to incorporate error or approximations into the modelling was critical in its implementation. For example, large uncertainty bounds covering an extensive range of feasible parameter values for the reverse direction of the esterase reaction were modelled. Despite the high levels of uncertainty for these parameters, the model is still able to make a good predictions, giving low *χ*^2^ values compared to simple models. This is particularly so for the complete reaction where error in these parameters have little impact on the prediction for the final alcohol concentration (Figures 5C, 6 and 7).

Furthermore, a number of approximations were necessary while modelling the ATP regeneration for the CAR step, resulting in a large uncertainty in the CAR reaction (Figure 3E). However high uncertainty in the CAR step did not necessarily result in high uncertainty in the complete reaction (Figure 5C and Figure 7), although some parameters had a larger effect once enzyme concentrations were minimized in the optimized reaction (Figure 6 and Figure 7). Sensitivity analysis suggests a more accurate characterization of the inhibition by PP_I_ on the CAR enzyme, as well as a better characterization of the AK reaction, would be good targets for reducing the uncertainty in the optimized reaction in future experiments.

### Conclusions

Multi-enzyme cascades, many of which are made possible via CAR enzymes, offer the opportunity to make an extended range of biocatalytic transformations economically viable. Here we show that thorough characterization of every component of a multi-enzyme cascade allows the development of a deterministic cascade model. This deterministic model robustly predicts the behavior of the cascade *in vitro* (*χ*^2^ = 3.0), demonstrating that such models are readily tractable for cascades of at least seven enzymes, a considerable advance on the most complex cascade previously described. We further demonstrate the utility of this model in understanding the reaction and for rapidly identifying steps in the catalysis that do not perform as expected. Finally, we exploited the model to optimize the transformation for maximum pathway flux with minimal enzyme usage. Our findings validate the use of deterministic models for *in vitro* biocatalysis, and strongly suggest that these should be used more widely in process development to increase the economic efficiency of enzyme cascades.

## Methods

### Materials and plasmids

All chemicals were purchased from Sigma-Aldrich (Gillingham, UK), and were of the highest purity available. Plasmid files are available as supplementary file 2. pNIC28-Bsa4 (68) is available through Addgene. Other pET vectors used in this study are available from Novagen. All reactions were performed in aqueous solutions. Methyl 4-toluate (#259667), 4-toluic acid (#T36803), 4-tolualdehyde (#35602) and 4-tolylalcohol (#127809) were each prepared at stock concentration of 500 mM in DMSO.

### Enzyme preparation

Genes for the expression of mpCAR, PTDH, tnAK, tnPAP, and ttPPiase were cloned into the *Nco*I and *Hind*III sites of pNIC28-Bsa4(68). The native *apADH* gene sequence was cloned previously into pET-30 Xa/LIC (69). The gene for mpCAR was obtained by PCR from genomic DNA. All other genes were codon optimized for *E. coli* and gene synthesized by IDT. All contained a N-terminal 6× histidine tag (68) (sequences available in Supplementary Information). Vectors were transformed into BL21 (DE3) *E. coli* for expression. mpCAR was co-transformed with a pCDF-Duet1 vector containing a phosphopantetheine transferase from *Bacillus subtilis*. A pET24-d plasmid containing afEst2 transformed into the *E. coli* strain BL21-CodonPlus (DE3)-RIPL (Agilent) has been described previously (42). Expression was carried out in LB media with the addition of 50 μg/ml of appropriate antibiotics.

Cells were grown to approximately 0.6 OD_600nm_ at 37 °C with shaking at 225 rpm, at which point IPTG was added to a concentration of 100 μM. Cells were then incubated overnight at 20 °C, except for afEst2 which was incubated overnight at 30 °C. Cells were harvested by centrifugation at 4,750 *g* and re-suspended in 25 mM Tris-HCl pH 8.0, 0.5 M NaCl_2_.

Cell lysate was prepared by sonication on ice followed by centrifugation at 20,000 *g* to remove the insoluble fraction. Enzymes were purified from the cell lysate using a 1 ml His-Trap FF crude column (GE Healthcare) using an elution gradient from 10 to 250 mM imidazole in 25 mM Tris-HCl pH 8.0, 0.5 M NaCl. The purified sample was then applied to a Superdex 200 HiLoad 16/60 gel filtration column (GE Healthcare) and eluted in 25 mM HEPES, pH 7.5, 0.1 M NaCl at 1.0 ml/min. Eluted fractions were analyzed by SDS-PAGE before being pooled and concentrated to between 1 and 8 mg/ml. To calculate protein concentration from OD_280nm_, an extinction coefficient and molecular weight for each enzyme was calculated using the ExPaSy ProtParam tool (70). Single use aliquots of protein were stored at −80 °C.

### Enzyme assays

Unless otherwise stated, all reactions were performed in triplicate in a 96-well microtiter plate using a Tecan M200 Infinite plate reader. All assays were carried out in a standard reaction buffer consisting of 100 mM Tris-HCl at pH 7.5, titrated to the correct pH whilst at 30 °C. All experiments used three experimental replicates, defined as experiments set up and run independently for each condition tested. Triplicate samples were sufficient to determine the necessary constants, whilst permitting sufficient throughput in single experiments.

### Buffers for pH vs activity assays

Buffers to measure activity across a range of pH values were prepared at 0.2 pH unit intervals at assay temperature. Buffers used were: 50 mM citrate between pH 4.0 and 5.8, 50 mM MES between pH 5.8 and 6.6, 50 mM PIPES between pH 6.4 and 7.4, 50 mM MOPS between pH 6.8 and 7.8, 50 mM HEPES between pH 7.0 and 8.0, 50 mM Tris between pH 7.8 and 9.0, and 50 mM boric acid between pH 8.8 and 10. 10 M and 1 M HCl or NaOH was used to titrate the buffers to the correct pH as appropriate.

### Thermostability assays

Thermostability was measured by heating enzyme samples for 30 min using the temperature gradient of a Bio-Rad thermocycler between 30 and 90 °C (or less where appropriate) before cooling on ice. Activity was then measured using specific enzyme assays (below) relative to a control sample kept on ice. Where appropriate, data were adapted from previous work as indicated in the relevant sections.

### CAR assays

To determine the two substrate kinetics of mpCAR with ATP and 4-toluic acid, reactions were set up containing 11 μg / ml mpCAR enzyme, 0.25 mM NADPH, 10 mM MgCl_2_, with a range of ATP and 4-toluic acid concentrations. The oxidation of NADPH was used to monitor reactions by measuring absorbance of NADPH at OD_340 nm_. Reactions were carried out for 10 minutes after a 5 minute pre-incubation at 30 °C. The appropriate equation was determined by fitting the initial rates of reaction at eight ATP concentrations around the expected *K*_*M*_ including a blank, each at five 4-toluic acid concentrations around its expected *K*_*M*_. Data were fitted by least squares non-linear regression using GraphPad Prism 5.0, and possible reaction mechanisms compared for goodness of fit. A *K*_*M*_ for NADPH was determined by setting up reactions containing 10 μg / ml mpCAR enzyme, 10 mM MgCl_2_, 1 mM ATP, 10 mM 4-toluic acid and a range of eight NADPH concentrations around its expected *K*_*M*_, after confirming NADPH concentration at OD_340 nm_. Other characterization was performed previously (15).

### ADH assays

To determine kinetic parameters for apADH, assays were carried out in sealed PCR tubes using a Bio-Rad thermocycler. After 15 to 30 minutes reactions were quenched by transfer into a 96-well microtiter plate containing 5 mM NaOH. Increase or decrease in NADH concentration was measured at OD_340nm_ to determine activity, relative to a blank reaction. Reactions to measure activity in the reductive direction were set up containing either 0.5 mM NADH or 10 mM 4-tolualdehyde. Reactions to measure activity in the oxidative direction were set up containing either 2.5 mM NAD^+^ or 100 mM 4-tolyl alcohol. In each case a range of concentrations were used for the other respective substrate around its expected *K*_*M*_, including a blank. 4-tolylaldehyde kinetics were carried out at 30 °C in standard reaction buffer using 166.7 μg / ml apADH enzyme. NAD^+^, NADH and 4-tolyl alcohol kinetics were carried out at 70 °C in 100 mM Tris-HCl titrated to pH 7.5 at 70 °C, using 40 μg / ml apADH enzyme. Only *K*_*M*_ constants were taken from data at 70 °C. Data were fitted by least squares non-linear regression using GraphPad Prism 5.0. Data for the effects of pH and temperature were adapted from previous work (46).

### PTDH assays

To determine kinetic parameters for PTDH, assays were set up containing 20 mM Na_2_HPO_3_, 2.8 μg / ml PTDH enzyme, and a range of NAD^+^ or NADP^+^ concentrations around the expected *K*_*M*_. The production of NADH or NADPH was monitored at OD_340nm_ and data were fitted by least squares non-linear regression using GraphPad Prism 5.0. The thermostability of a sample containing 18 μg / ml PTDH was measured as described above. Activity at various pH values was determined using a range of buffers described previously, in place of the standard reaction buffer.

### PPIase

PPiase activity was measured using the production of phosphomolybdate to measure phosphate content, as described (71). Using a standard curve, the rate of phosphate production by ttPPiase at 0.22 μg / ml was determined using a high (5 mM) concentration of PP_I_ in the presence of 10 mM MgCl_2_, in standard reaction buffer. Five measurements were taken in triplicate over 20 minutes. A conservative *K*_*M*_ was estimated from the BRENDA database (72) and the rate of PP_I_ hydrolysis used to calculate *k*_*cat*_.

Activity at pH was determined using the same phosphomolybdate method with reaction buffers covering a range of pH values, detailed above. The thermostability of a sample containing 0.44 μg / ml ttPPiase was measured using the assay described previously, with readings taken at time points 0 and 20 minutes. 10 mM MgCl_2_ was included in the heated sample.

### PAP assays

ADP formation was measured using an ADP-Glo kinase assay kit from Promega in a 384-well solid white microtiter plate, following the manufacturer’s instructions. Assays were carried out in standard reaction buffer containing 20 or 40 mM MgCl_2_, 12.5 mM polyphosphate, 3.25 mM AMP, 12.5 μg / ml or 125 μg / ml of tnPAP enzyme (as indicated in SI). Polyphosphate was calculated as concentration of Na.PO_3_ units, with a molecular weight of 102. Reactions were carried out for 15 minutes before quenching with the ADP-Glo kit. A range of polyphosphate, AMP and MgCl_2_ concentrations around an expected *K*_*M*_ were assayed in turn. ADP production was calculated from a standard curve, and the data fitted by least squares non-linear regression using GraphPad Prism 5.0. Specific assay conditions are shown in Supplementary Figures 35 to 38. Activity at different pH values was measured using a range of buffers described above in place of the standard reaction buffer. Thermostability was determined as described above, using the ADP-Glo assay to measure residual relative activity.

### Mathematical modelling

Mathematical modelling was carried out in Python 3.4 using the SciPy (73) and SALib (74) modules. A python package (https://github.com/willfinnigan/kinetics) was developed during the course of this work. Outputs were exported to GraphPad Prism 5.0 for plotting. The integrate.odeint function in SciPy was used to solve ordinary differential equations.

Uncertainty analysis was carried out making use of the SALib module (74). Input bounds for each parameter were defined as either the calculated 95 % confidence intervals or where uncertainty was judged to be high 25 or 50 % of the parameter value. Sampling of the possible inputs within these bounds was carried out by Latin hypercube sampling of 1,000 samples, with the model for each sample run. At each time point the mean, 5^th^ and 95^th^ percentile of each substrate concentration was plotted to represent the uncertainty of the model.

Reduced chi-squared values were calculated using the formula (48):

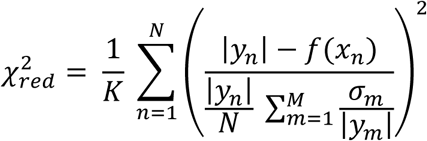

Where *K* is the number of degrees of freedom; *N* is the total number of experimental points determined; *M* is the total number of experimental points determined for each substrate/intermediate/product; |*y*_*m/n*_| is the mean experimental value at each point determined; *f(x*_*n*_) is the model derived value for y at time x; and *s*_*n*_ is the standard deviation in each experimental value. The reduced chi-squared formula was modified here to take the mean fractional standard deviation for each chemical: this approach was taken to mitigate the effect of (random) tightly determined triplicate points dominating the calculation. Three models were used for each dataset: the median deterministic model described above; a quadratic equation of the form y = ax^2^ + bx + c fitted to the experimental data; and a linear equation of the form y = ax + c fitted to the experimental data.

The method of Sobol (75) was used for sensitivity analysis as part of the SALib module (74), although numerus methods are available. Second order effects were not calculated, and the total sensitivity indices plotted to show the sensitivity of each parameter. The sample number was set at 1,000 (chosen for consistency with past studies; this was sufficient to obtain a good sampling rate whilst not overloading computing resources). For more information on the use of uncertainty and sensitivity analysis in process modelling please see the following references (76,77).

### Optimization using a genetic algorithm

A custom built, single objective genetic algorithm was used to minimize enzyme cost, on the condition over 90 % alcohol yield was reached. A general outline of the algorithm is shown in Supplementary Figure 44.

### Model validation reactions

Standard curves of methyl 4-toluate, 4-toluic acid, 4-tolylaldehyde and 4-tolyl alcohol were prepared and analyzed by HPLC. 500 μl reactions were set up in triplicate in 1.5 ml microcentrifuge tubes, and incubated in a thermoshaker (EpMotion T5075t thermal module, Eppendorf) at 30 °C with 500 rpm shaking. 50 μl samples were taken every 30 minutes, mixed with 50 μl acetonitrile, and centrifuged for 10 minutes. Supernatant was removed and stored at 4 °C before being analyzed by HPLC.

### HPLC

An Eclipse Plus C18 column (Agilent) with a particle size of 3.5 μm, measuring 4.6 × 100 mm, was used. The column was run at 60 °C on the following method using two buffers: buffer A: 95 % H_2_O, 5 % (v/v) acetonitrile, 0.1 % (v/v) trifluoroacetic acid; buffer B: 5 % H_2_0, 95 % (v/v) acetonitrile, 0.1 % (v/v) trifluoroacetic acid. 3 μl of sample was injected and eluted on a gradient from 0 to 100 % buffer B over 10 minutes. Buffer B was maintained at 100 % for a further 2 minutes before the column was re-equilibrated with buffer A for 2 minutes.

### Mass spectrometry

Low and high resolution mass spectra were obtained by staff at the University of Manchester. Electrospray (ES) spectra were recorded on a Waters Platform II with an SQ Detector 2. High resolution mass spectra (HRMS) were recorded on a Thermo Finnigan MAT95XP and are accurate to ± 0.001 Da.

## Acknowledgements

The authors thank Andrew Hill (University of Manchester) for providing the pCDF-Sfp plasmid. We also thank Colin Edge, Dan Tray and Neil Hodnett (GlaxoSmithKline) for their advice on process modelling; and Dr Khulood Alyahya (University of Exeter) for advice on model quality. Additionally we thank Anna Peters (University of Manchester) for her assistance with the mass spectrometry. WF was funded by BBSRC (grant no. BB/K501001/1) and GlaxoSmithKline. RC was funded by BBSRC and GlaxoSmithKline.

## References

1. Pollard DJ, Woodley JM. Biocatalysis for pharmaceutical intermediates: the future is now. Trends Biotechnol. 2007;25(2):66–73.

2. Turner NJ, Truppo MD. Biocatalysis enters a new era. Curr Opin Chem Biol. Elsevier Ltd; 2013;17(2):212–4.

3. Oroz-Guinea I, García-Junceda E. Enzyme catalysed tandem reactions. Curr Opin Chem Biol. 2013;17(2):236–49.

4. Wells A, Meyer HP. Biocatalysis as a strategic green technology for the chemical industry. ChemCatChem. 2014;6(4):918–20.

5. Wilding KM, Schinn SM, Long EA, Bundy BC. The emerging impact of cell-free chemical biosynthesis. Curr Opin Biotechnol. Elsevier Ltd; 2018;53(1):115–21.

6. du Preez R, Clarke KG, Callanan LH, Burton SG. Modelling of immobilised enzyme biocatalytic membrane reactor performance. J Mol Catal B Enzym. Elsevier B.V.; 2015;119:48–53.

7. Bujara M, Schümperli M, Billerbeck S, Heinemann M, Panke S. Exploiting cell-free systems: Implementation and debugging of a system of biotransformations. Biotechnol Bioeng. 2010;106(3):376–89.

8. Hold C, Panke S. Towards the engineering of in vitro systems. J R Soc Interface. 2009;6(Suppl_4):S507–21.

9. Xue R, Woodley JM. Process technology for multi-enzymatic reaction systems. Bioresour Technol. Elsevier Ltd; 2012;115:183–95.

10. Winkler M. Carboxylic acid reductase enzymes (CARs). Curr Opin Chem Biol. Elsevier Ltd; 2018;43(Figure 2):23–9.

11. Qu G, Guo J, Yang D, Sun Z. Biocatalysis of carboxylic acid reductases: Phylogenesis, catalytic mechanism and potential applications. Green Chem. Royal Society of Chemistry; 2018;20(4):777–92.

12. Turner NJ, O’Reilly E. Biocatalytic retrosynthesis. Nat Chem Biol. Nature Publishing Group; 2013;9(5):285–8.

13. France SP, Hussain S, Hill AM, Hepworth LJ, Howard RM, Mulholland KR, et al. One-Pot Cascade Synthesis of Mono- and Disubstituted Piperidines and Pyrrolidines using Carboxylic Acid Reductase (CAR), ω-Transaminase (ω-TA), and Imine Reductase (IRED) Biocatalysts. ACS Catal. 2016;6(6):3753–9.

14. Khusnutdinova AN, Flick R, Popovic A, Brown G, Tchigvintsev A, Nocek B, et al. Exploring Bacterial Carboxylate Reductases for the Reduction of Bifunctional Carboxylic Acids. Biotechnol J. 2017;12(11):1–12.

15. Finnigan W, Thomas A, Cromar H, Gough B, Snajdrova R, Adams JP, et al. Characterization of Carboxylic Acid Reductases as Enzymes in the Toolbox for Synthetic Chemistry. ChemCatChem. 2017;9(6):1005–17.

16. Hansen EH, Møller BL, Kock GR, Bünner CM, Kristensen C, Jensen OR, et al. De novo biosynthesis of Vanillin in fission yeast (schizosaccharomyces pombe) and baker’s yeast (saccharomyces cerevisiae). Appl Environ Microbiol. 2009;75(9):2765–74.

17. Kallio P, Pásztor A, Thiel K, Akhtar MK, Jones PR. An engineered pathway for the biosynthesis of renewable propane. Nat Commun. 2014;5:4731.

18. Zhang Y-HP. Production of biofuels and biochemicals by in vitro synthetic biosystems: Opportunities and challenges. Biotechnol Adv. 2015;33(7):1467–83.

19. Kunjapur AM, Cervantes B, Prather KLJ. Coupling carboxylic acid reductase to inorganic pyrophosphatase enhances cell-free in vitro aldehyde biosynthesis. Biochem Eng J. Elsevier B.V.; 2016;109:19–27.

20. Ni Y, Holtmann D, Hollmann F. How green is biocatalysis? to calculate is to know. ChemCatChem. 2014;6(4):930–43.

21. Woodyer R, Van der Donk WA, Zhao H. Relaxing the nicotinamide cofactor specificity of phosphite dehydrogenase by rational design. Biochemistry. 2003;42(40):11604–14.

22. McLachlan MJ, Johannes TW, Zhao H. Further Improvement of Phosphite Dehydrogenase Thermostabilty by Saturation Mutagenesis. Biotechnol Bioeng. 2007;99(2):268–74.

23. Zhao H, Van Der Donk WA. Regeneration of cofactors for use in biocatalysis. Curr Opin Biotechnol. 2003;14(6):583–9.

24. Woodyer RD, Johannes TW, Zhao H. Regeneration of cofactors for use in biocatalysis. In: Enzyme Technology. New Delhi, India.: Asiatech Publishers Inc.,; 2003. p. 85–103.

25. Kameda A, Shiba T, Kawazoe Y, Satoh Y, Ihara Y, Munekata M, et al. A novel ATP regeneration system using polyphosphate-AMP phosohotransferase and polyphosphate kinase. J Biosci Bioeng. 2001;91(6):557–63.

26. Resnick SM, Zehnder AJB. In vitro ATP regeneration from polyphosphate and AMP by polyphosphate:AMP phosphotransferase and adenylate kinase from Acinetobacter johnsonii 210A. Appl Environ Microbiol. 2000;66(5):2045–51.

27. Medema MH, van Raaphorst R, Takano E, Breitling R. Computational tools for the synthetic design of biochemical pathways. Nat Rev Microbiol. Nature Publishing Group; 2012;10(3):191–202.

28. Kuhn D, Blank LM, Schmid A, Bühler B. Systems biotechnology - Rational whole-cell biocatalyst and bioprocess design. Eng Life Sci. 2010 Oct 7;10(5):384–97.

29. Ellis T, Adie T, Baldwin GS. DNA assembly for synthetic biology: from parts to pathways and beyond. Integr Biol (Camb). 2011 Feb;3(2):109–18.

30. Purcell O, Jain B, Karr JR, Covert MW, Lu TK. Towards a whole-cell modeling approach for synthetic biology. Chaos. 2013;23(2).

31. Vasić-Rački D, Findrik Z, Vrsalović Presečki A. Modelling as a tool of enzyme reaction engineering for enzyme reactor development. Appl Microbiol Biotechnol. 2011;91(4):845–56.

32. Ringborg RH, Woodley JM. The application of reaction engineering to biocatalysis. React Chem Eng. Royal Society of Chemistry; 2016;1(1):10–22.

33. Sin G, Woodley JM, Gernaey K V. Application of modeling and simulation tools for the evaluation of biocatalytic processes: A future perspective. Biotechnol Prog. 2009;25(6):1529–38.

34. Rios-Solis L, Morris P, Grant C, Odeleye AOO, Hailes HC, Ward JM, et al. Modelling and optimisation of the one-pot, multi-enzymatic synthesis of chiral amino-alcohols based on microscale kinetic parameter determination. Chem Eng Sci. Elsevier; 2015;122(June 2014):360–72.

35. Peri S, Karra S, Lee YY, Karim MN. Modeling intrinsic kinetics of enzymatic cellulose hydrolysis. Biotechnol Prog. 2007;23(3):626–37.

36. Findrik Z, Vasić-Rački D. Biotransformation of D-Methionine into L-Methionine in the Cascase of Four Enzymes. Biotechnol Bioeng. 2007;98:956.

37. Yadav GD, Manjula Devi K. A kinetic model for the enzyme-catalyzed self-epoxidation of oleic acid. JAOCS, J Am Oil Chem Soc. 2001;78(4):347–51.

38. Sayar NA, Chen BH, Lye GJ, Woodley JM. Process modelling and simulation of a transketolase mediated reaction: Analysis of alternative modes of operation. Biochem Eng J. 2009;47(1–3):10–8.

39. Rollin JA, Martin del Campo J, Myung S, Sun F, You C, Bakovic A, et al. High-yield hydrogen production from biomass by in vitro metabolic engineering: Mixed sugars coutilization and kinetic modeling. Proc Natl Acad Sci. 2015;112(16):4964–9.

40. Schrittwieser JH, Sattler J, Resch V, Mutti FG, Kroutil W. Recent biocatalytic oxidation-reduction cascades. Curr Opin Chem Biol. 2011;15(2):249–56.

41. Guy JE, Isupov MN, Littlechild JA. The Structure of an Alcohol Dehydrogenase from the Hyperthermophilic Archaeon Aeropyrum pernix. J Mol Biol. 2003;2836(03):1041–51.

42. Sayer C, Finnigan W, Isupov MN, Levisson M, Kengen SWM, van der Oost J, et al. Structural and biochemical characterisation of Archaeoglobus fulgidus esterase reveals a bound CoA molecule in the vicinity of the active site. Sci Rep. Nature Publishing Group; 2016;6(1):25542.

43. Bonting CFC, Kortstee GJJ, Zehnder AJB. Properties of polyphosphate - AMP phosphotransferase of Acinetobacter strain-210A. J Bacteriol. 1991;173(20):6484–8.

44. Sato M, Masuda Y, Kirimura K, Kino K. Thermostable ATP regeneration system using polyphosphate kinase from Thermosynechococcus elongatus BP-1 for D-amino acid dipeptide synthesis. J Biosci Bioeng. 2007;103(2):179–84.

45. Vieille C, Krishnamurthy H, Hyun H-H, Savchenko A, Yan H, Zeikus JG. Thermotoga neapolitana adenylate kinase is highly active at 30 degrees C. Biochem J. 2003;372(Pt 2):577–85.

46. Hirakawa H, Kamiya N, Kawarabayashi Y, Nagamune T. Properties of an alcohol dehydrogenase from the hyperthermophilic archaeon Aeropyrum pernix K1. J Biosci Bioeng. 2004;97(3):202–6.

47. Cornish-Bowden A. Fundamentals of Enzyme Kinetics, 4th Edition. Wiley; 2012.

48. Andrae R, Schulze-Hartung T, Melchior P. Dos and don’ts of reduced chi-squared. 2010;1–12.

49. Kraut J. Serine proteases: Struture and mechanism of catalysis. Annu Rev Biochem. 1977;46:331–58.

50. Schellenberger V, Jakubke H -D. Protease-Catalyzed Kinetically Controlled Peptide Synthesis. Angew Chemie Int Ed English. 1991;30(11):1437–49.

51. Gaertner, H. & Pugserver, A. Kinetics and specifisity of serine proteases in peptide synthesis catalysed in organic solvents. Eur J Biochem. 1989;181:207–13.

52. Gahloth D, Dunstan MS, Quaglia D, Klumbys E, Lockhart-Cairns MP, Hill AM, et al. Structures of carboxylic acid reductase reveal domain dynamics underlying catalysis. Nat Chem Biol. 2017;13(9):975–81.

53. Cornish-Bowden A. Fundamentals of enzyme kinetics. 2004. 153–156 p.

54. Mu H, Zhou SM, Xia Y, Zou H, Meng F, Yan Y Bin. Inactivation and unfolding of the hyperthermophilic inorganic pyrophosphatase from Thermus thermophilus by sodium dodecyl sulfate. Int J Mol Sci. 2009;10(6):2849–59.

55. Mitic A, Skov T, Gernaey K V. Removal of benzaldehyde from a water/ethanol mixture by applying scavenging techniques. Green Process Synth. 2017;6(3):353–61.

56. Bubb WA, Berthon HA, Kuchel PW. Tris buffer reactivity with low-molecular-weight aldehydes: NMR characterization of the reactions of glyceraldehyde 3-phosphate. Bioorganic Chemistry. 1995. p. 119–30.

57. Lin JL, Palomec L, Wheeldon I. Design and analysis of enhanced catalysis in scaffolded multienzyme cascade reactions. ACS Catal. 2014;4(2):505–11.

58. Wheeldon I, Minteer SD, Banta S, Barton SC, Atanassov P, Sigman M. Substrate channelling as an approach to cascade reactions. Nat Chem. Nature Publishing Group; 2016;8(4):299–309.

59. Nossent J, Elsen P, Bauwens W. Sobol’ sensitivity analysis of a complex environmental model. Environ Model Softw. Elsevier Ltd; 2011;26(12):1515–25.

60. Tullman-Ercek D. Metabolism: “Channeling” Hans Krebs. Nat Chem Biol. Nature Publishing Group; 2015;11(3):180–1.

61. Zheng M, Zhang S, Ma G, Wang P. Effect of molecular mobility on coupled enzymatic reactions involving cofactor regeneration using nanoparticle-attached enzymes. J Biotechnol. Elsevier B.V.; 2011;154(4):274–80.

62. Akhtar MK, Turner NJ, Jones PR. Carboxylic acid reductase is a versatile enzyme for the conversion of fatty acids into fuels and chemical commodities. Proc Natl Acad Sci U S A. 2013 Jan 2;110(1):87–92.

63. Blair JMD. Magnesium, Potassium, and the Adenylate Kinase Equilibrium: Magnesium as a Feedback Signal from the Adenine Nucleotide Pool. Eur J Biochem. 1970;13(2):384–90.

64. Sheng XR, Li X, Pan XM. An iso-random Bi Bi mechanism for adenylate kinase. J Biol Chem. 1999;274(32):22238–42.

65. Vinnakota KC, Wu F, Kushmerick MJ, Beard DA. Chapter 2 Multiple Ion Binding Equilibria, Reaction Kinetics, and Thermodynamics in Dynamic Models of Biochemical Pathways. 1st ed. Methods in Enzymology. Elesvier Inc.; 2009. 29–68 p.

66. Deb K, Agrawal S, Pratab A, Meyarivan T. A Fast Elitist Non-Dominated Sorting Genetic Algorithm for Multi-Objective Optimization: NSGA-II. Proc 6^{th} Conf Parallel Probl Solving from Nat. 2000;849–58.

67. Tufvesson P, Lima-Ramos J, Nordblad M, Woodley JM. Guidelines and cost analysis for catalyst production in biocatalytic processes. Org Process Res Dev. 2011;15(1):266–74.

68. Savitsky P, Bray J, Cooper CDO, Marsden BD, Mahajan P, Burgess-brown NA, et al. High-throughput production of human proteins for crystallization: The SGC experience. J Struct Biol. Elsevier Inc.; 2010;172(1):3–13.

69. Hickey AM. Unpublished work. 2008.

70. Gasteiger E, Hoogland C, Gattiker A, Duvaud S, Wilkins M., R.D A, et al. Protein Identification and Analysis Tools on the ExPASy Server. John M. Walker (ed): The Proteomics Protocols Handbook, Humana Press (2005). 2005. p. 571–607.

71. Heinonen JK, Lahti RJ. A new and convenient colorimetric determination of inorganic orthophosphate and its application to the assay of inorganic pyrophosphatase. Anal Biochem. 1981;113(2):313–7.

72. Schomburg I, Chang A, Placzek S, Sohngen C, Rother M, Lang M, et al. BRENDA in 2013: integrated reactions, kinetic data, enzyme function data, improved disease classification: new options and contents in BRENDA. Nucleic Acids Res. 2013;41:764–72.

73. Jones E, Oliphant T, Peterson P. SciPy: Open source scientific tools for Python. 2001.

74. Herman J, Usher W. SALib: An open-source Python library for Sensitivity Analysis. J Open Source Softw. 2017;2(9):10–1.

75. Sobol IM. Global sensitivity indices for nonlinear mathematical models and their Monte Carlo estimates. Math Comput Simul. 2001;55:271–80.

76. Sin G, Gernaey K V, Lantz AE. Good modelling practice (GMoP) for PAT applications: Propagation of input uncertainty and sensitivity analysis. Biotechnol Prog. 2009;25:1043–53.

77. Price JA, Mathias N, Woodley J, Kjobsted J. Application of Uncertainty and Sensitivity Analysis to a Kinetic Model for Enzymatic Biodiesel Production. Proc 12th IFAC Symp Comput Appl Biotechnol. 2013;12(1):161–8.

78. Itoh H, Shiba T. Polyphosphate synthetic activity of polyphosphate:AMP phosphotransferase in Acinetobacter johnsonii 210A. J Bacteriol. 2004;186(15):5178–81.

79. Perrier V, Burlacu-Miron S, Bourgeois S, Surewicz WK, Gilles a M. Genetically engineered zinc-chelating adenylate kinase from Escherichia coli with enhanced thermal stability. J Biol Chem. 1998;273(30):19097–101.

