## Supplementary figures for "Engineering a seven enzyme biotransformation using mathematical modelling and characterized enzyme parts"

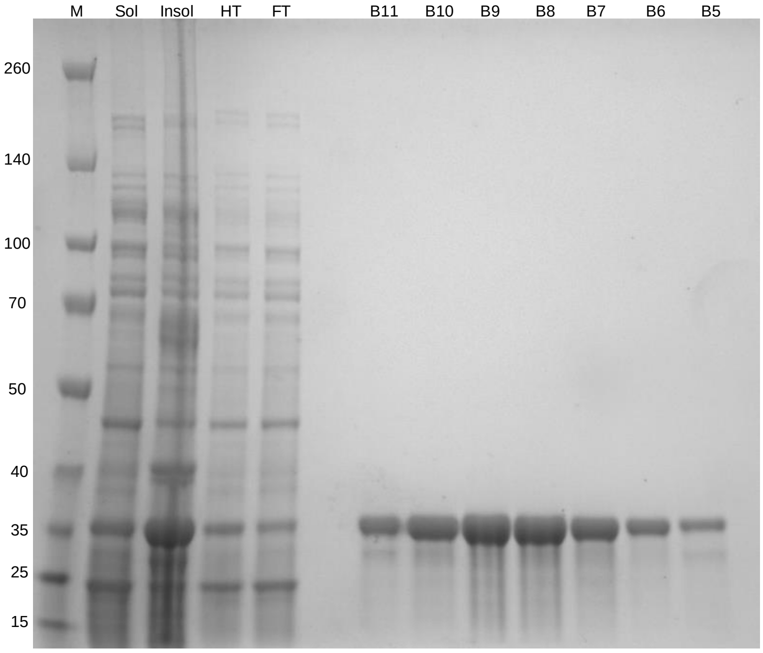


**Supplementary Figure 1 - SDS-PAGE analysis of the AF-Est2 protein purification**
Sol: Soluble fraction of cell lysate. Insol: Insoluble fraction of cell lysate. HT: Soluble lysate heat treated at 70 °C for 30 minutes before removal of precipitated proteins by centrifugation. FT: The flow through after loading the heat treated sample onto the nickel column. B11-B5: Fractions collected from the gel filtration column believed to be the purified AF-Est2 protein with an expected MW of 29 kDa.

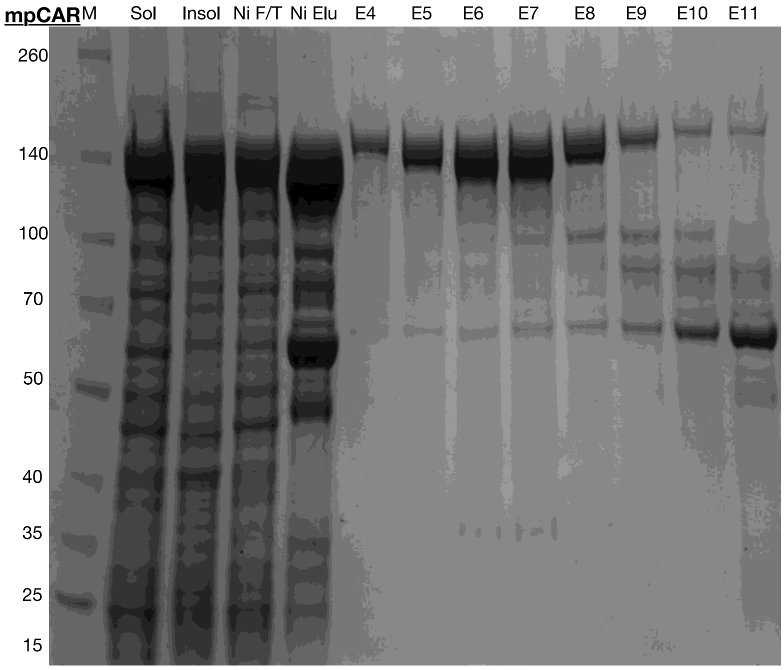


**Supplementary Figure 2 - SDS-PAGE analysis of the mpCAR protein purification.**
Sol: Soluble fraction of cell lysate. Insol: Insoluble fraction of cell lysate. Ni F/T: The flow through after loading the sample onto the nickel column. Ni Elu: Purified protein after the nickel purification step. E4-E11: Fractions collected from the gel filtration column believed to be the purified mpCAR protein with an

expected MW of 128 kDa. Fractions E9 – E11 contained another unknown protein at approximately 60 kDa. These fractions were not pooled with the remaining fractions.

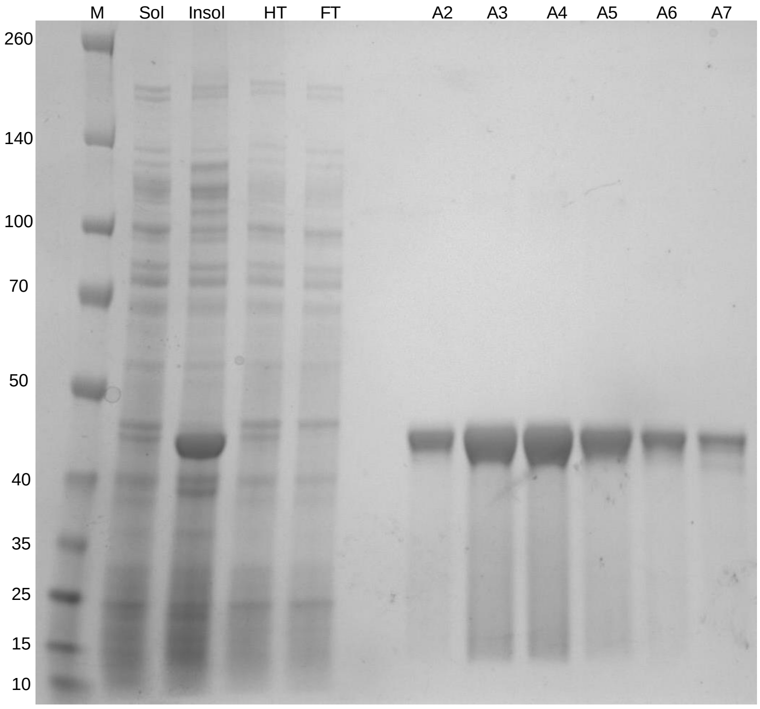


**Supplementary Figure 3 - SDS-PAGE analysis of the ApADH protein purification.**
Sol: Soluble fraction of cell lysate. Insol: Insoluble fraction of cell lysate. HT: Soluble lysate heat treated at 70 °C for 30 minutes before removal of precipitated proteins by centrifugation. FT: The flow through after loading the heat treated sample onto the nickel column. A2-A7: Fractions collected from the gel filtration column believed to be the purified ApADH protein with an expected MW of 41 kDa.

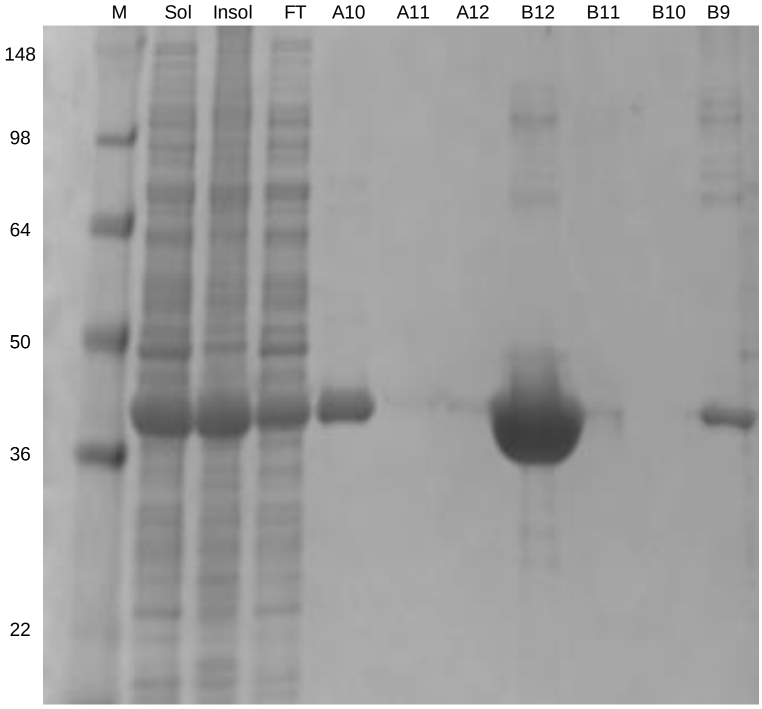


**Supplementary Figure 4 - SDS-PAGE analysis of the PTDH protein purification.**
Sol: Soluble fraction of cell lysate. Insol: Insoluble fraction of cell lysate. FT: The flow through after loading the sample onto the nickel column. A10-B9: Fractions collected from the gel filtration column believed to be the purified PTDH protein with an expected MW of 38.7 kDa. Very little protein was

detected by SDS-PAGE in lanes A11, A112, B11 and B10, however there was likely a problem in running in the gel as there was an unavoidable delay between preparing the samples and running them. Samples at the extremes of the collected peak show pure protein so the entire peak was used.

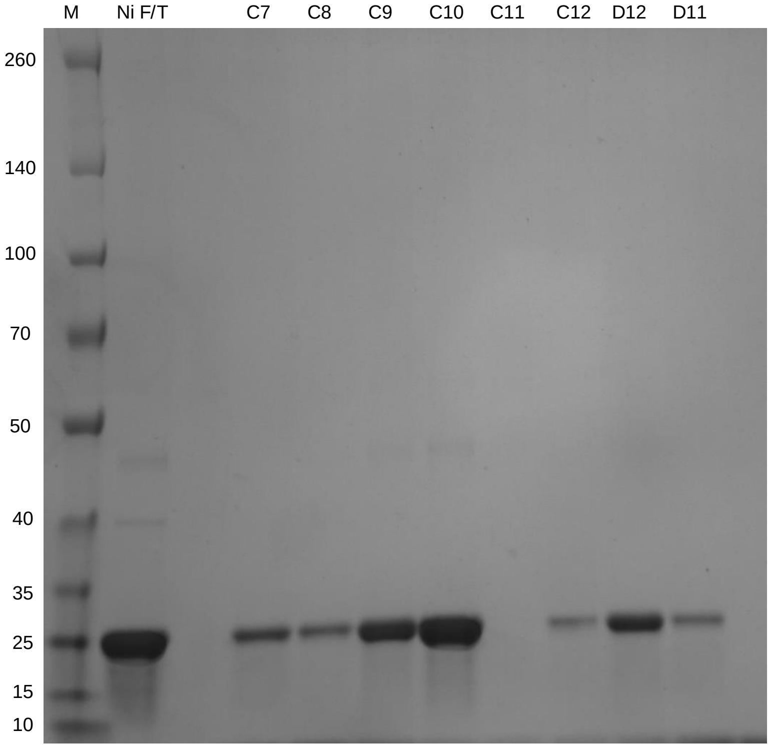


**Supplementary Figure 5 - SDS-PAGE analysis of the ttPpiase protein purification.**
Ni F/T: The flow through after loading the heat treated sample onto the nickel column. C7-D11: Fractions collected from the gel filtration column believed to be the purified ttPpiase protein with an expected MW of 21.8 kDa.

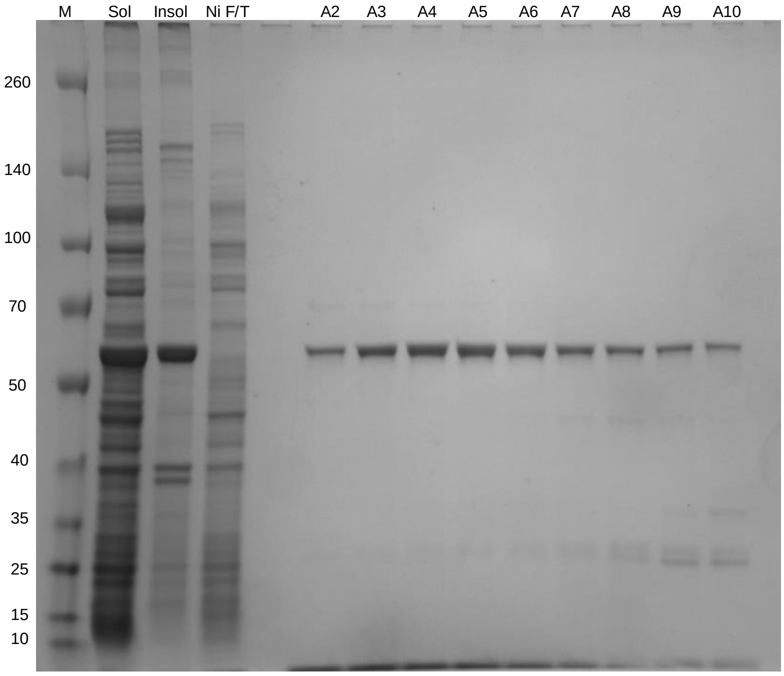


**Supplementary Figure 6 - SDS-PAGE analysis of the tnPAP protein purification.**
Sol: Soluble fraction of cell lysate. Insol: Insoluble fraction of cell lysate. Ni F/T: The flow through after loading the sample onto the nickel column. A2-A10: Fractions collected from the gel filtration column believed to be the purified tnPAP protein with an expected MW of 61.1 kDa.

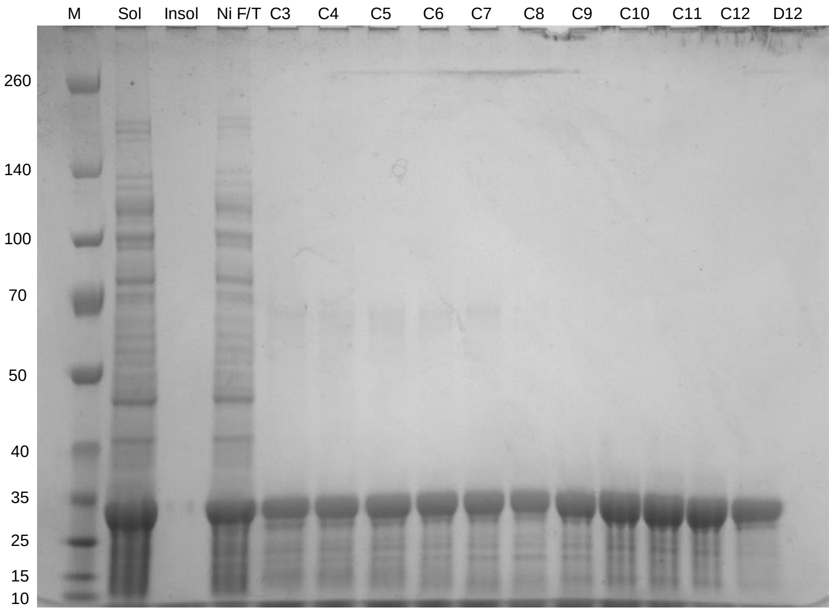


**Supplementary Figure 7 - SDS-PAGE analysis of the tnAK protein purification.**
Sol: Soluble fraction of cell lysate. Insol: Insoluble fraction of cell lysate. Ni F/T: The flow through after loading the sample onto the nickel column. C3-D12: Fractions collected from the gel filtration column believed to be the purified tnPPT protein with an expected MW of 27.6 kDa.

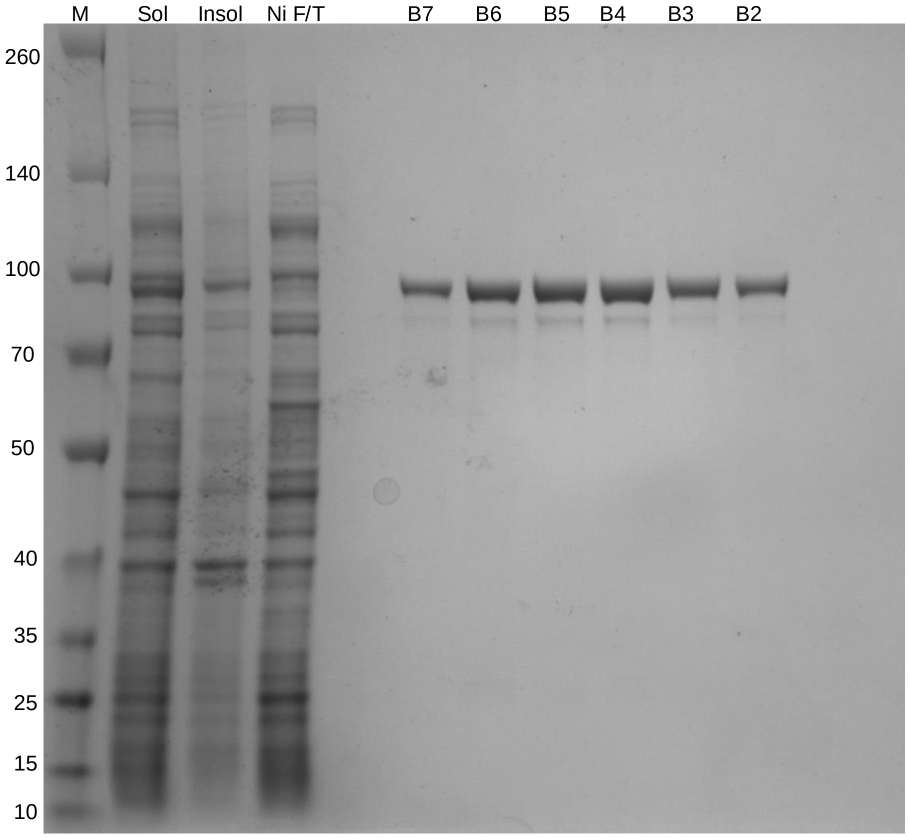


**Supplementary Figure 8 - SDS-PAGE analysis of the tePPK protein purification.**
Sol: Soluble fraction of cell lysate. Insol: Insoluble fraction of cell lysate. Ni F/T: The flow through after loading the sample onto the nickel column. B7-B2: Fractions collected from the gel filtration column believed to be the purified tePPK protein with an expected MW of 87.5 kDa.

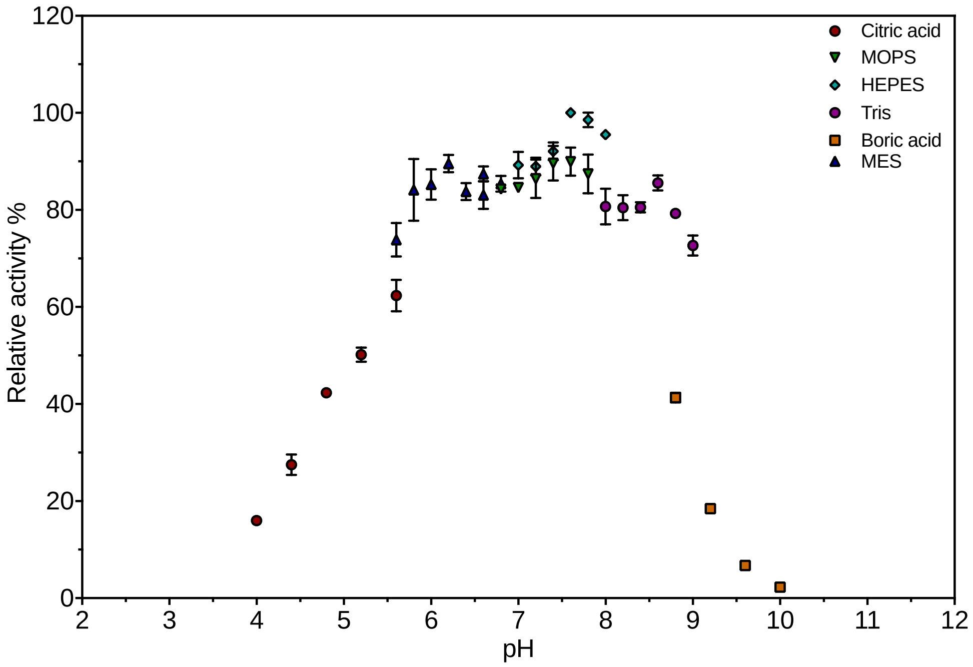


**Supplementary Figure 9 – PTDH activity at different pH values**

Activity of PTDH at various pH values, relative to the maximum activity detected. Various buffers were used to cover the range of pH values as indicated.

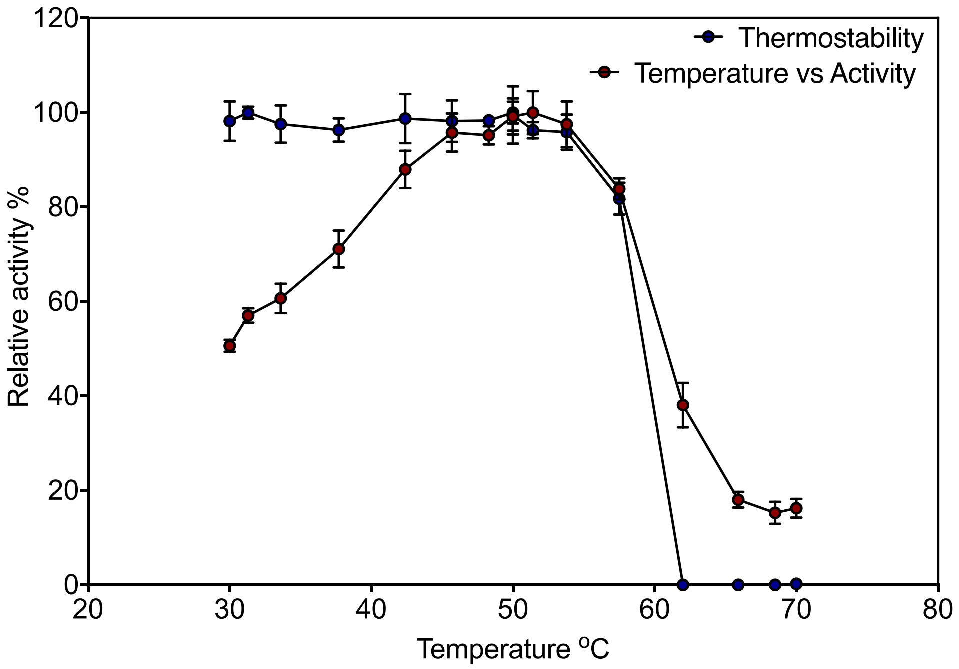


**Supplementary Figure 10 – PTDH thermostability and temperature vs activity**
Red circles show PTDH activity at temperature, relative to the maximum activity at 51.4 °C. Blue circles show residual activity after incubating PTDH at various temperatures for 30 minutes, relative to a control kept on ice.

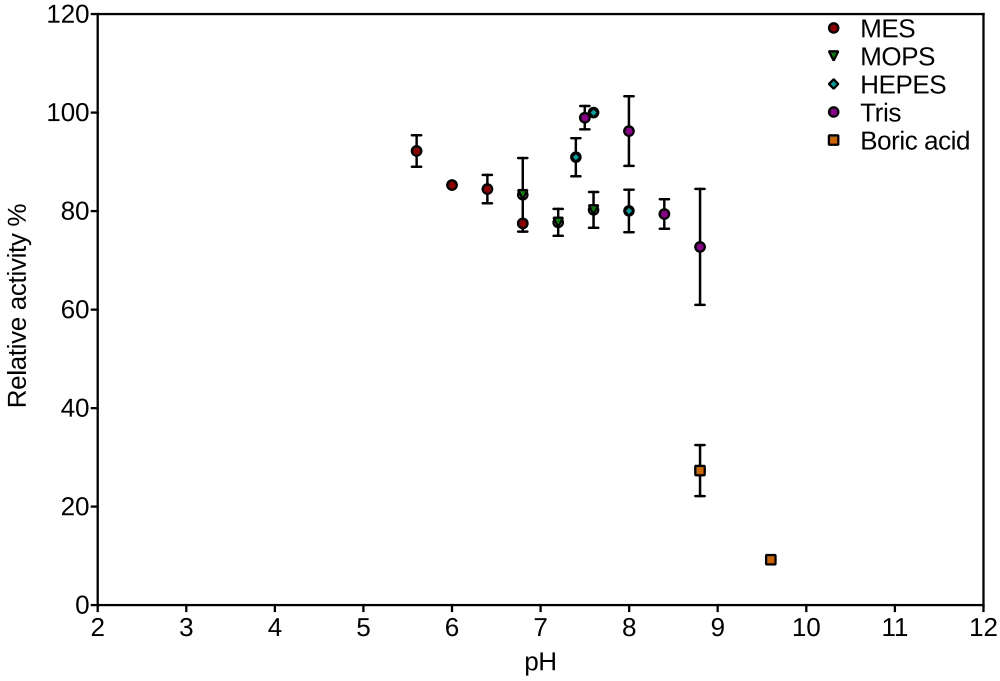


**Supplementary Figure 11 – tnPAP activity at different pH values**
Activity of tnPAP at various pH values, relative to the maximum activity detected. Various buffers were used to cover the range of pH values as indicated.

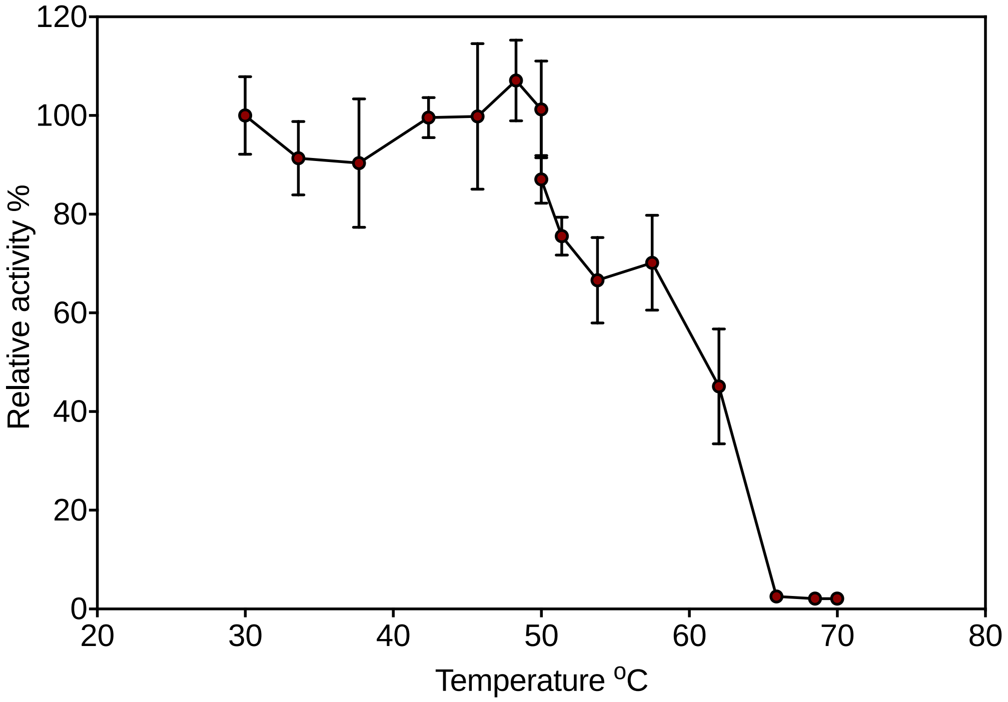


**Supplementary Figure 12 – tnPAP thermostability**

Residual activity after incubating tnPAP at various temperatures for 30 minutes, relative to a control kept on ice.

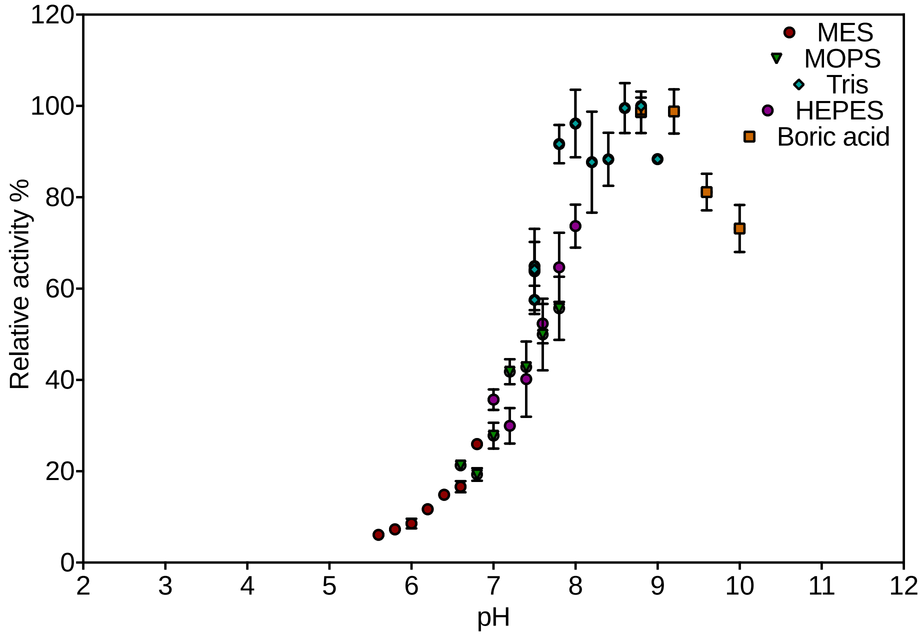


**Supplementary Figure 13 – ttPPiase activity at pH**

Activity of ttPPiase at various pH values, relative to the maximum activity detected. Various buffers were used to cover the range of pH values as indicated.

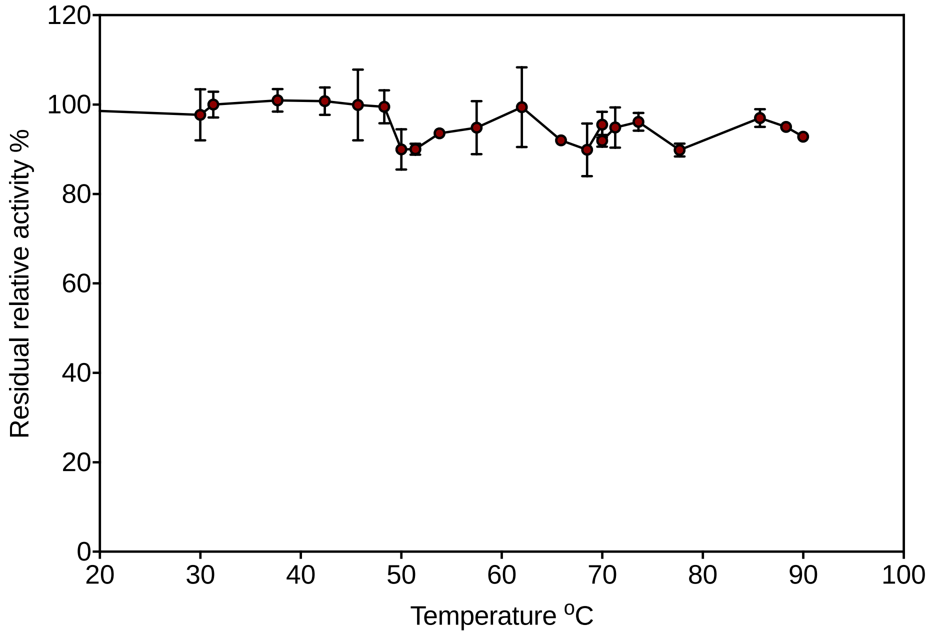


**Supplementary Figure 14 - ttPPiase thermostability**
Residual activity after incubating ttPPiase at various temperatures for 30 minutes, relative to a control kept on ice.

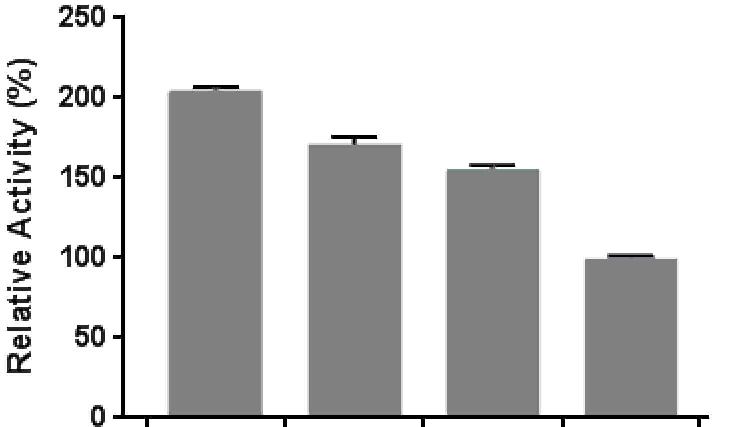


| | | | | |
|---|---|---|---|---|
| 100 | 2.5 | 100 | 2.5 | Tris concentration (mM) |
| + | + | - | - | Auxiliary reagents |
| 203.9 | 170.7 | 155.1 | 100 | Mean (%) |

| | F Value. | P Value. |
|---|---|---|
| Tris - HCl. | 166.17 | 3.8e-11 |
| Auxiliary reagents. | 319.01 | 9.3e-14 |
| Tris - HCl:Addtives. | 23.62 | 9.5e-5 |

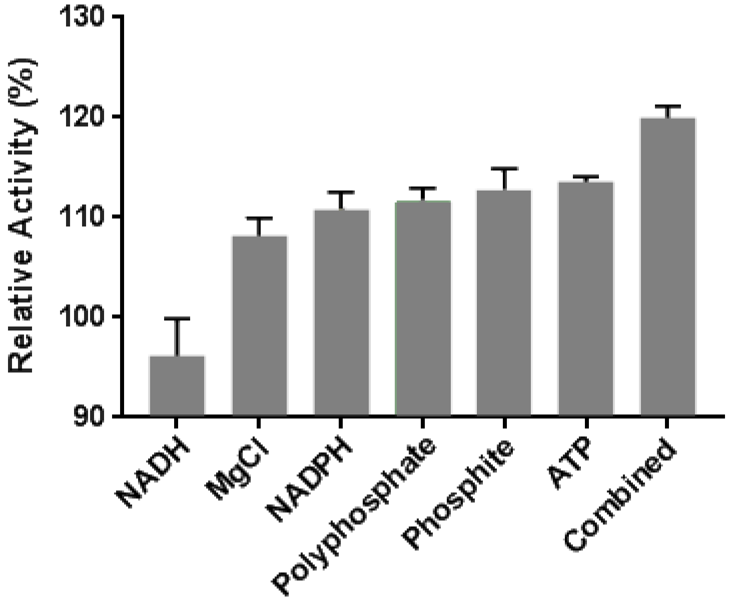


**Supplementary Figure 15 – The rate of afEst2 increases upon the addition of final reaction components.** The afEST2 enzyme assay (Sayer et al., 2017) was performed in conditions replicating the differences between initial assay and final assay conditions. The reaction was initiated by the addition of 500 µM of substrate (4-nitrophenyl-butyrate). The production of 4-nitrophenol was observed at 405 nm. A. Effect of auxiliary reactants and buffer. The reaction was performed in the presence of 2.5 or 100 mM Tris-HCl pH 7.5, and with or without the auxiliary reactants from the final assay (NADH (500 µM), MgCl2 (20,000 µM), NADPH (500 µM), polyphosphate (6,000 µM), phosphite (20,000 µM) or ATP (1,250 µM)). An increase in buffer concentration causes a 55% increase in rate (p < 10-10), whilst the addition of auxiliary reagents causes a 70% increase in rate (p < 10-13). The addition of the auxiliary reagents to the high buffer condition causes a further increase of ~30%, implying that there is a less than additive effect of the two changes (p < 10-4; all statistics calculated using a two-way ANOVA with Tukey's post-hoc determination). Experiments were performed in sextuplet. B. Effects of individual auxiliary reactants. The reaction was performed in the presence of each auxiliary reactant alone and compared to control, in 100 mM Tris-HCl pH 7.5. The afEST2 rate increased compared to the control for all of the auxiliary reactants tested, with the exception of NADH. The sum of these increases was considerably more than the increase seen for the combination of all reagents, implying that the effects of these are not necessarily specific. This further implies that most likely the effect of these treatments is ionic stabilization of the protein rather than a specific interaction. Experiments were performed in triplicate. Error bars show SEM.

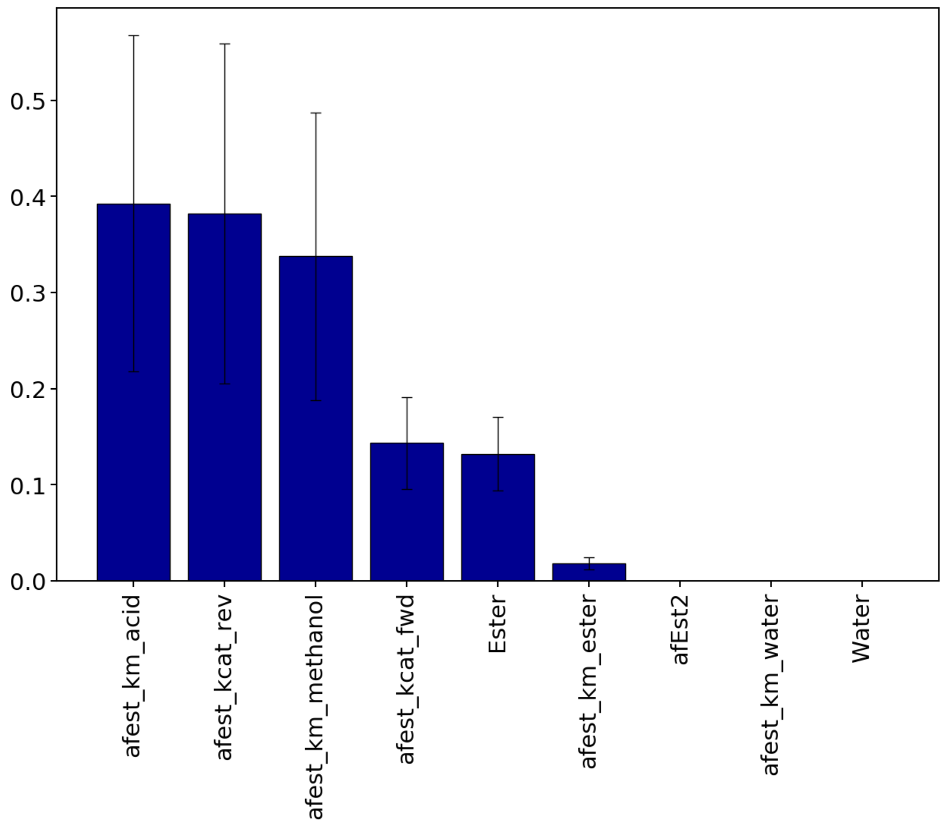


**Supplementary Figure 16 - Sensitivity analysis of the reversible afEst2 reaction.**
The total sensitivity indices (ST) are shown which take into account 1st order and all other interactions. Sensitivity is in reference to the uncertainty in the final methyl-*p*-toluate concentration. Error bars show the 95 % confidence intervals. The sum of all sensitivity indices' should equal 1.

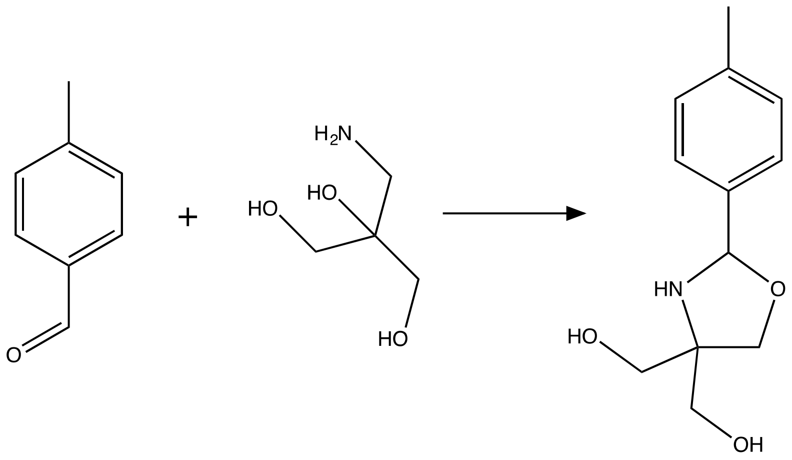


**Supplementary Figure 17 – Proposed side reaction between 4-methylbenzalde and Tris**
The proposed side reaction yields a product with an exact mass of 223.1.

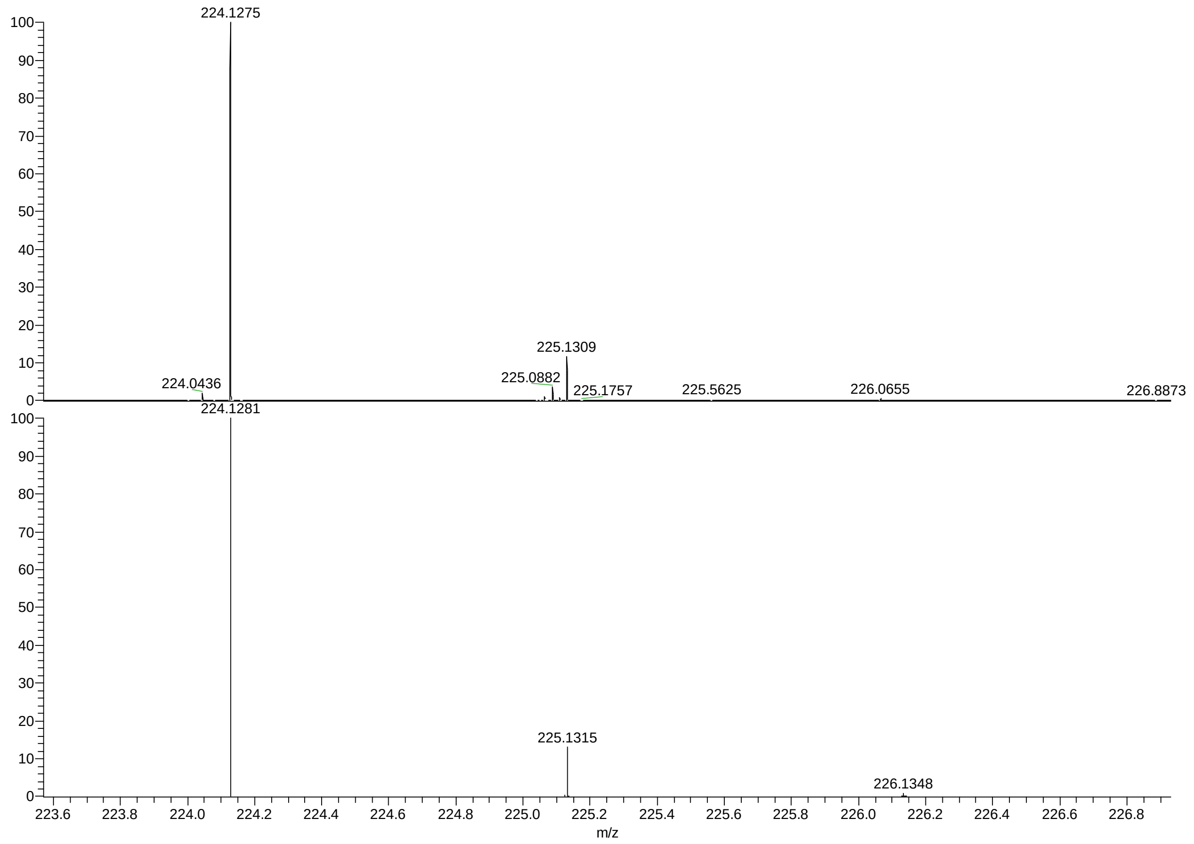


**Supplementary Figure 18 – Mass spec analysis of a reaction between 4 mM 4-methylbenzaldehyde and 100 mM Tris.**

The recorded HRMS and the theoretical isotopic pattern for the product proposed in Figure 5 – Supplementary Figure 1. 4 mM 4-tolualdehyde was incubated in 100 mM Tris-HCl pH 7.5 overnight at room temperature, and a sample taken. The recorded data and the theoretical isotopic pattern for the proposed and the produced products are in accordance with each other.

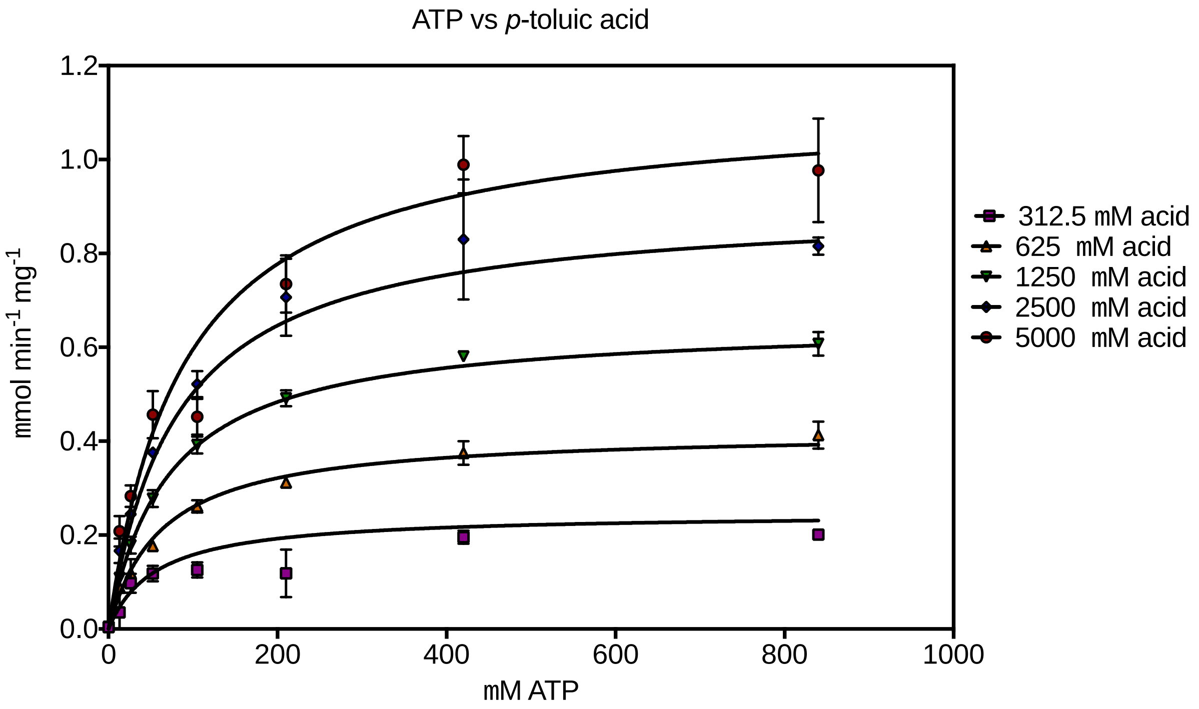


**Supplementary Figure 19 - The kinetics of mpCAR with varying concentrations of ATP and 4-toluic acid.** Data were fitted best to the equation for a steady-state sequential reaction.

**Supplementary Figure 20 – Parameters calculated fitting kinetics of mpCAR with varying concentrations of ATP and 4-toluic acid to a sequential steady state equation.**

$$v = V_{max} \cdot \frac{[\mathrm{A}]\cdot[\mathrm{B}]}{(K_I^A \cdot K_M^B)+(K_M^A\cdot[\mathrm{B}])+(K_M^B\cdot[\mathrm{A}])+([\mathrm{A}]\cdot[\mathrm{B}])}$$

| **Best-fit values** | |
|---|---|
| $V_{MAX}$ (µmol / min / mg) | 1.5 |
| $K_I$ ATP (µM) | 50 |
| $K_M$ Acid (µM) | 1,500 |
| $K_M$ ATP (µM) | 100 |
| **Std. Error** | |
| $V_{MAX}$ (µmol / min / mg) | 0.07 |
| $K_I$ ATP (µM) | 20 |
| $K_M$ Acid (µM) | 200 |
| $K_M$ ATP (µM) | 10 |

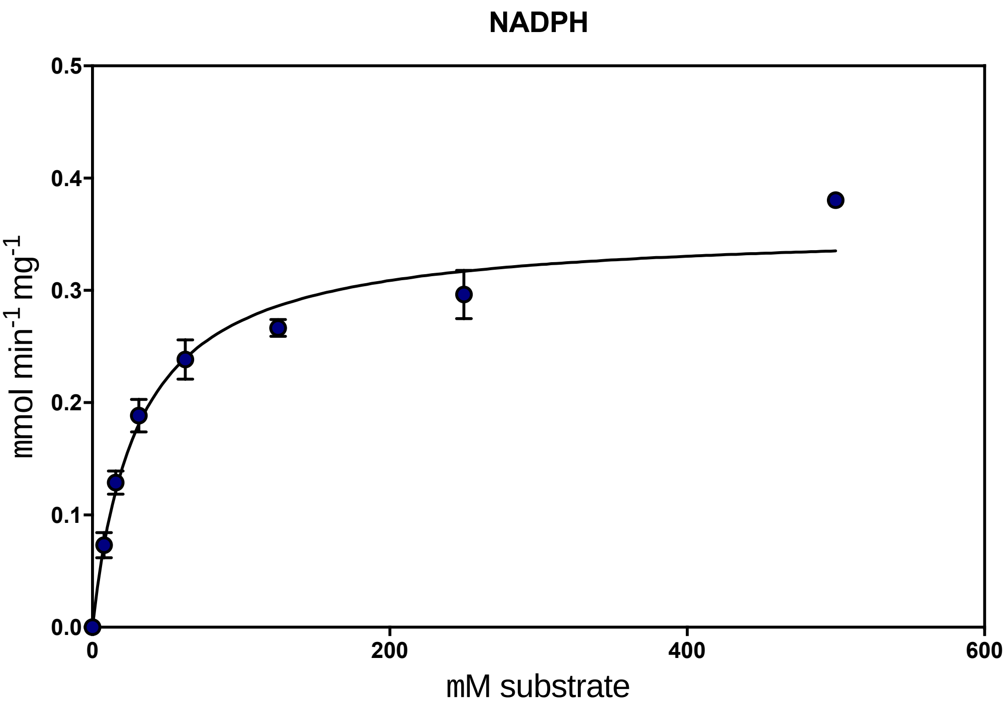


**Supplementary Figure 21 – The kinetics of mpCAR with varying concentrations NADPH.** Data have been fitted to the Michaelis-Menten equation.

| **Supplementary Figure 22 -Parameters calculated fitting kinetics of mpCAR with varying concentrations of NADPH to the Michaelis-Menten equation.** | |
|---|---|
| **Best-fit values** | |
| $V_{MAX}$ (µmol / min / mg) | 0.35 |
| $K_M$ NADPH (µM) | 30 |
| **Std. Error** | |
| $V_{MAX}$ (µmol / min / mg) | 0.01 |
| $K_M$ NADPH (µM) | 4 |

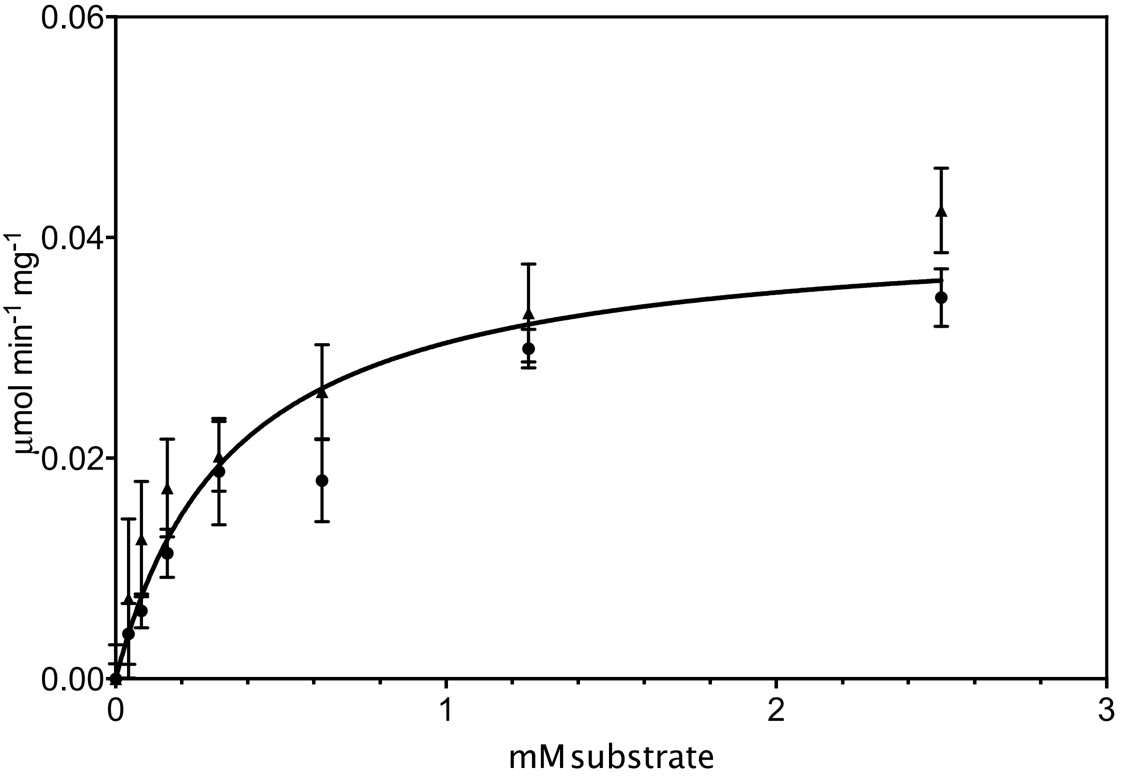


**Supplementary Figure 23 - The kinetics of apADH in the reductive direction at 30 °C, pH 7.5 with varying concentrations of 4-tolualdehyde.**

Two experiments were carried out shown as triangles and circles, with the data combined for fitting to the Michaelis-Menten equation.

| **Supplementary Figure 24 - Parameters calculated fitting kinetics of apADH in the reductive direction at 30 °C, pH 7.5, with varying concentrations of 4-tolualdehyde to the Michaelis-Menten equation.** | |
|---|---|
| **Best-fit values** | |
| $V_{MAX}$ (µmol / min / mg) | 0.041 |
| $K_M$ *p*-tolualdehyde (µM) | 350 |
| **Std. Error** | |
| $V_{MAX}$ (µmol / min / mg) | 0.002 |
| $K_M$ *p*-tolualdehyde (µM) | 60 |

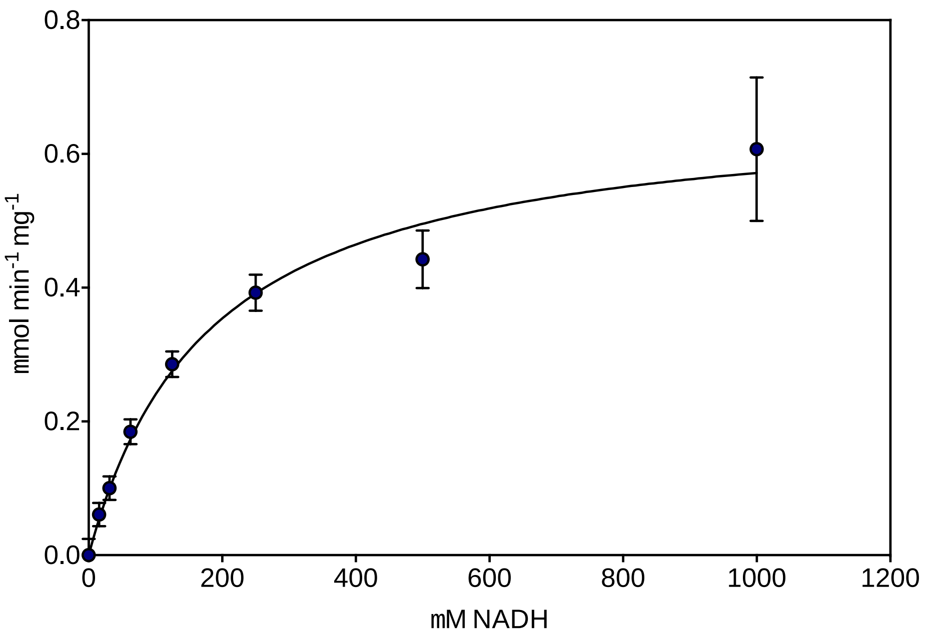


**Supplementary Figure 25 - The kinetics of apADH in the reductive direction at 70 °C, pH 7.5 with varying concentrations of NADH.**
Data have been fitted to the Michaelis-Menten equation.

| **Supplementary Figure 26 - Parameters calculated fitting kinetics of apADH in the reductive direction at 70 °C, pH 7.5, with varying concentrations of NADH, to the Michaelis-Menten equation.** Only $K_M$ used in this work. | |
|---|---|
| **Best-fit values** | |
| $V_{MAX}$ (µmol / min / mg) | 0.68 |
| $K_M$ NADH (µM) | 180 |
| **Std. Error** | |
| $V_{MAX}$ (µmol / min / mg) | 0.04 |
| $K_M$ NADH (µM) | 30 |

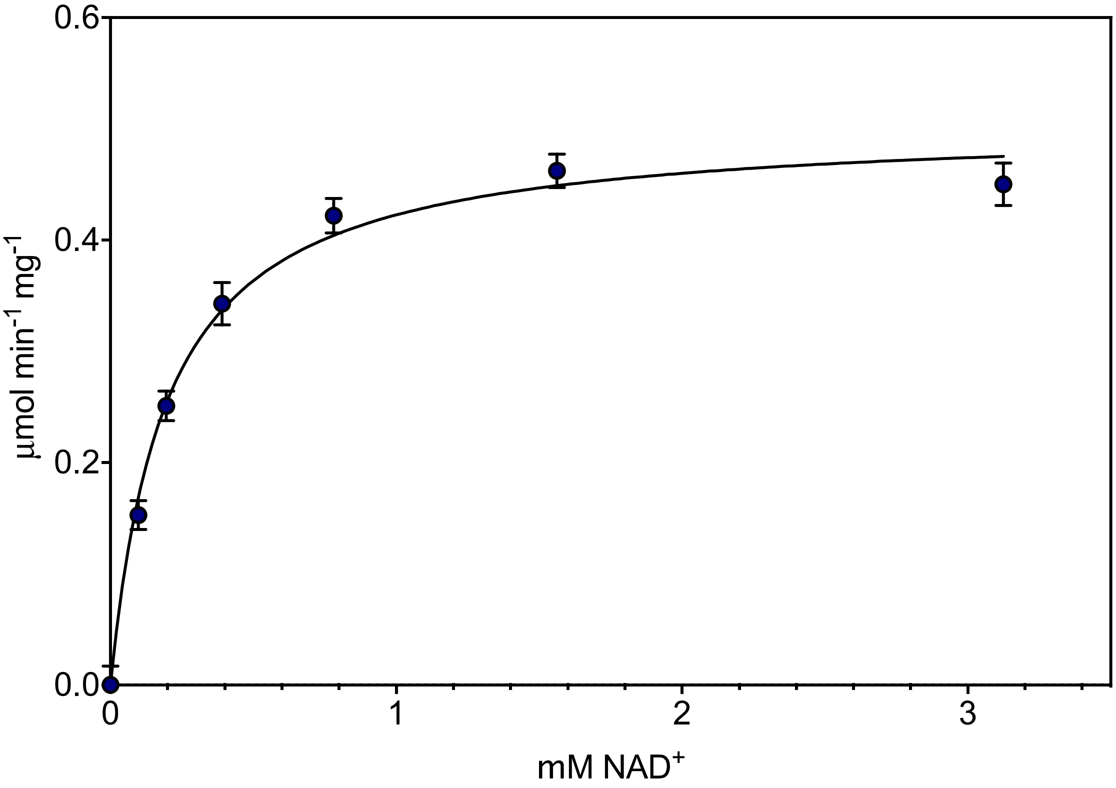


**Supplementary Figure 27 - The kinetics of apADH in the oxidative direction at 70 °C, pH 7.5 with varying concentrations of $NAD^+$.**
Data have been fitted to the Michaelis-Menten equation.

**Supplementary Figure 28 - Parameters calculated fitting kinetics of apADH in the oxidative direction at 70 °C, pH 7.5, with varying concentrations of $NAD^+$, to the Michaelis-Menten equation.**
Only $K_M$ used in this work.

| | |
|---|---|
| **Best-fit values** | |
| $V_{MAX}$ (µmol / min / mg) | 0.50 |
| $K_M$ $NAD^+$ (µM) | 195 |
| **Std. Error** | |
| $V_{MAX}$ (µmol / min / mg) | 0.01 |
| $K_M$ $NAD^+$ (µM) | 16 |

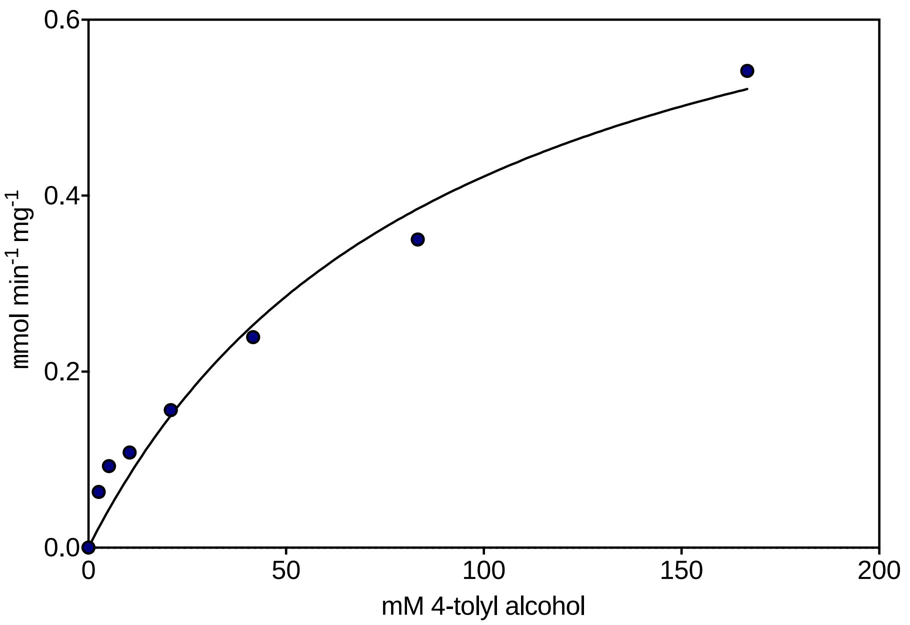


**Supplementary Figure 29 - The kinetics of apADH in the oxidative direction at 70 °C, pH 7.5 with varying concentrations of 4-tolyl alcohol.**
Data have been fitted to the Michaelis-Menten equation.

**Supplementary Figure 30 - Parameters calculated fitting kinetics of apADH in the oxidative direction at 70 °C, pH 7.5, with varying concentrations of $NAD^+$, to the Michaelis-Menten equation.**
As substrate concentration could not be taken high enough to calculate an accurate $K_M$ or $V_{MAX}$ this data was not used. An approximate $K_M$ of 100 mM was used and kcat in the oxidative direction assumed to be approximately equal to the forward.

| | |
|---|---|
| **Best-fit values** | |
| $V_{MAX}$ (µmol / min / mg) | 0.80 |
| $K_M$ *p*-tolyl alcohol (mM) | 90 |
| **Std. Error** | |
| $V_{MAX}$ (µmol / min / mg) | 0.06 |
| $K_M$ *p*-tolyl alcohol (mM) | 15 |

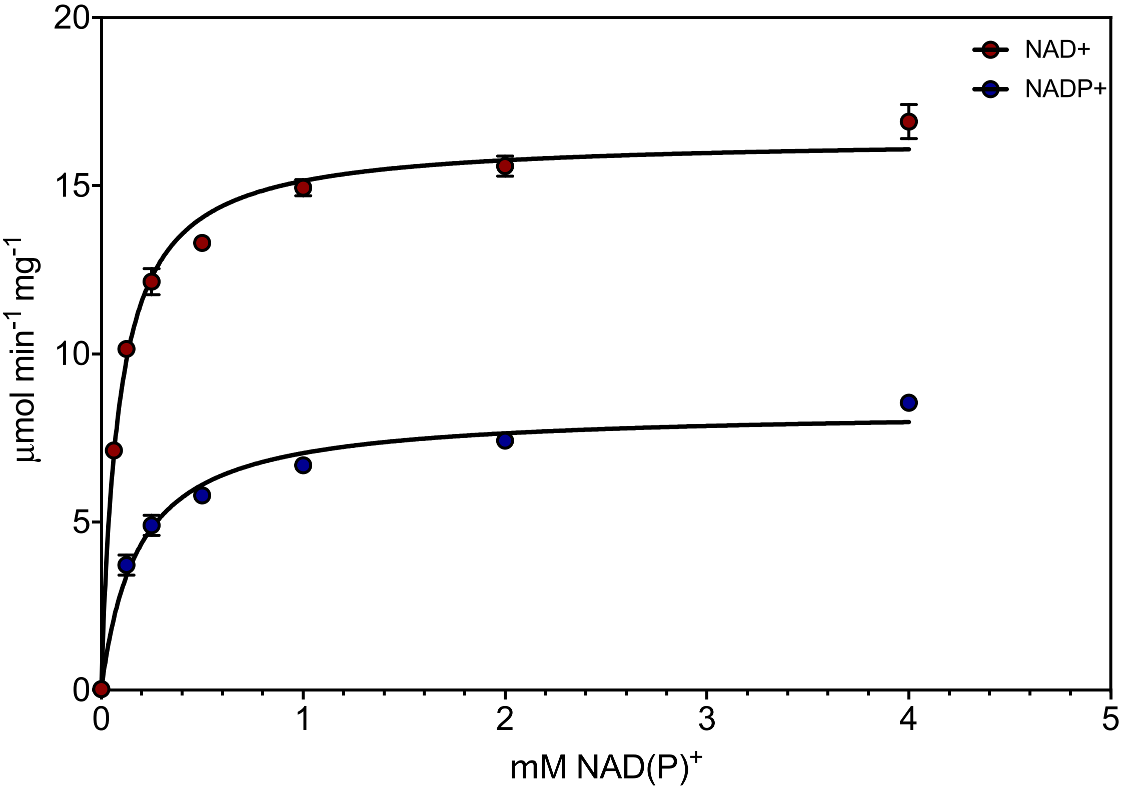


**Supplementary Figure 31 – The kinetics of PTDH with varying concentrations of NADP$^+$ and NAD$^+$.** Data have been fitted to the Michaelis-Menten equation.

**Supplementary Figure 32 - Parameters calculated fitting kinetics of PTDH with varying concentrations of NADP$^+$ and NAD$^+$ to the Michaelis-Menten equation.**

| **Best-fit values** | **NADP$^+$** | **NAD$^+$** |
|---|---|---|
| $V_{MAX}$ (µmol / min / mg) | 8.3 | 16.4 |
| $K_M$ NAD(P)$^+$( µM) | 180 | 85 |
| **Std. Error** | | |
| $V_{MAX}$ (µmol / min / mg) | 0.2 | 0.2 |
| $K_M$ 4-tolyl alcohol (mM) | 20 | 5 |

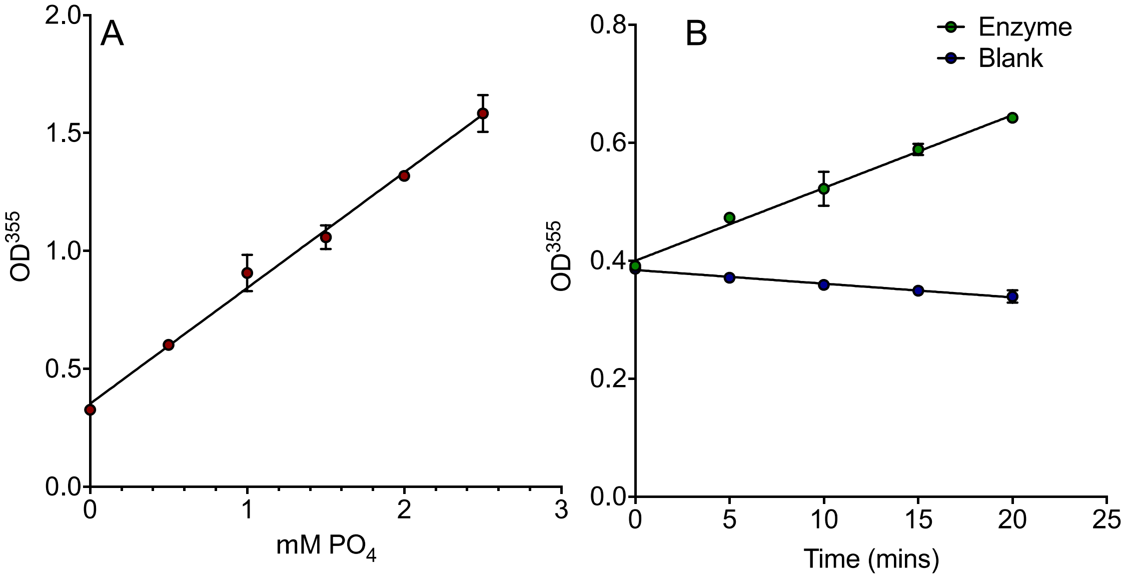


**Supplementary Figure 33 – Determining *$k_{cat}$* for ttPPiase**

A: A standard curve of phosphate concentration vs OD 335 nm was determined for the assay.

B: The rate of phosphate production was determined (green circles), along with a blank rate (blue circles). Rate of blank subtracted phosphate production was calculated from the standard curve, from which *$k_{cat}$* was calculated.

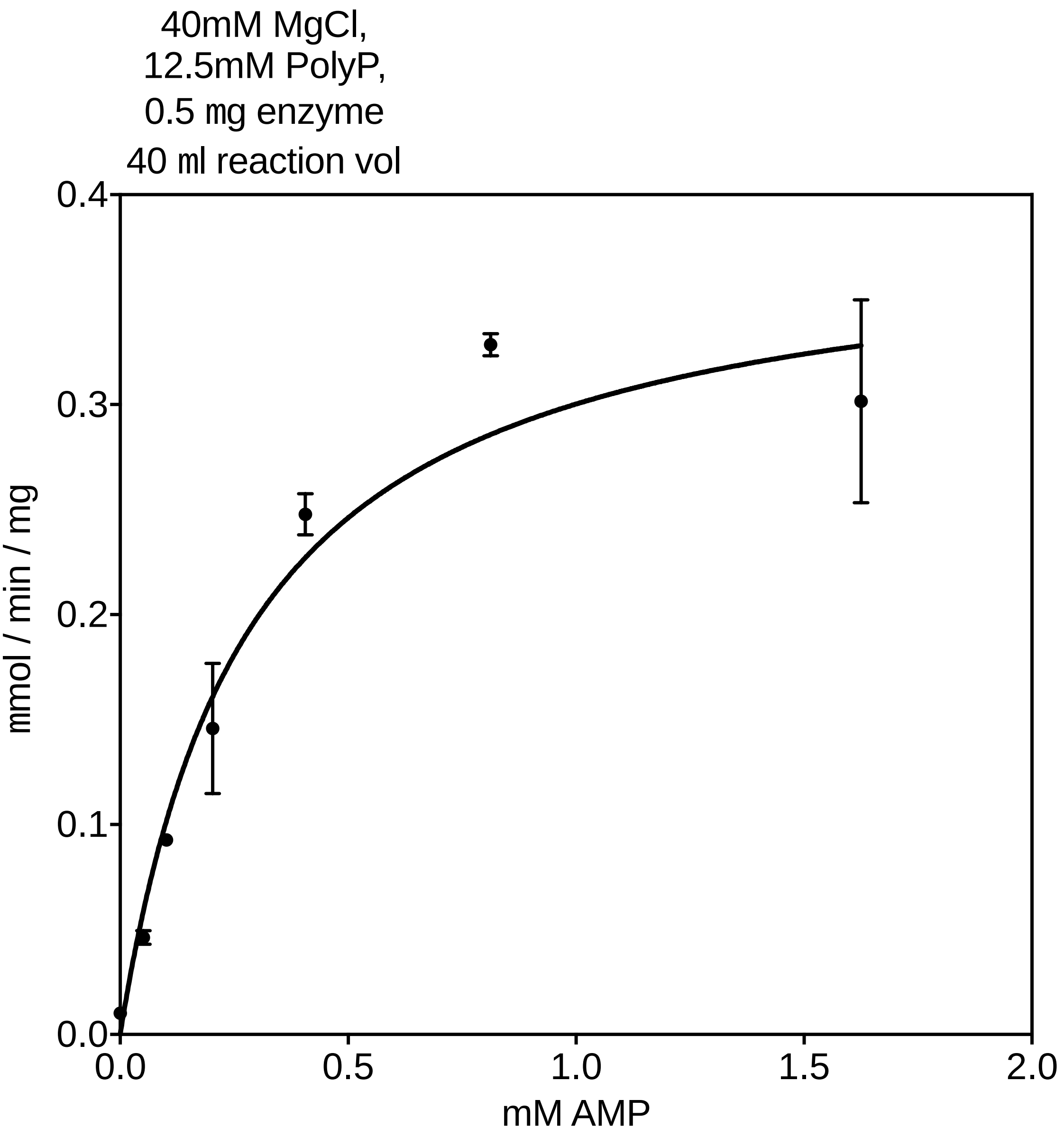


**Supplementary Figure 34 – The kinetics of tnPAP with varying concentrations of AMP**
Data fitted to the Michaelis-Menten equation.

| **Supplementary Figure 35 - Parameters calculated fitting kinetics of tnPAP with varying concentrations of AMP to the Michaelis-Menten equation.** | |
|---|---|
| **Best-fit values** | |
| $V_{MAX}$ (µmol / min / mg) | 0.39 |
| $K_M$ AMP (mM) | 0.28 |
| **Std. Error** | |
| $V_{MAX}$ (µmol / min / mg) | 0.03 |
| $K_M$ AMP (mM) | 0.05 |

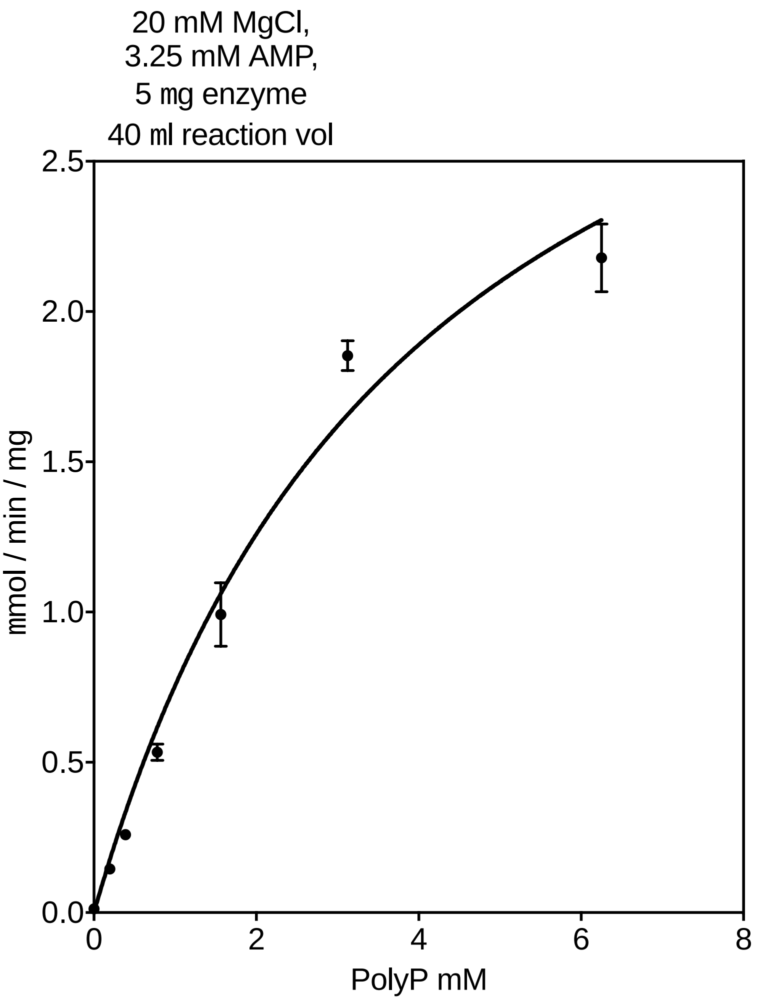


**Supplementary Figure 36 - The kinetics of tnPAP with varying concentrations of Polyphosphate (PolyP)**
Data have been fitted to the Michaelis-Menten equation.

**Supplementary Figure 37 - Parameters calculated fitting kinetics of tnPAP with varying concentrations of polyphosphate to the Michaelis-Menten equation.**

| | |
|---|---|
| Best-fit values | |
| Vmax | 3.781 |
| Km | 4.004 |
| Std. Error | |
| Vmax | 0.3454 |
| Km | 0.6808 |

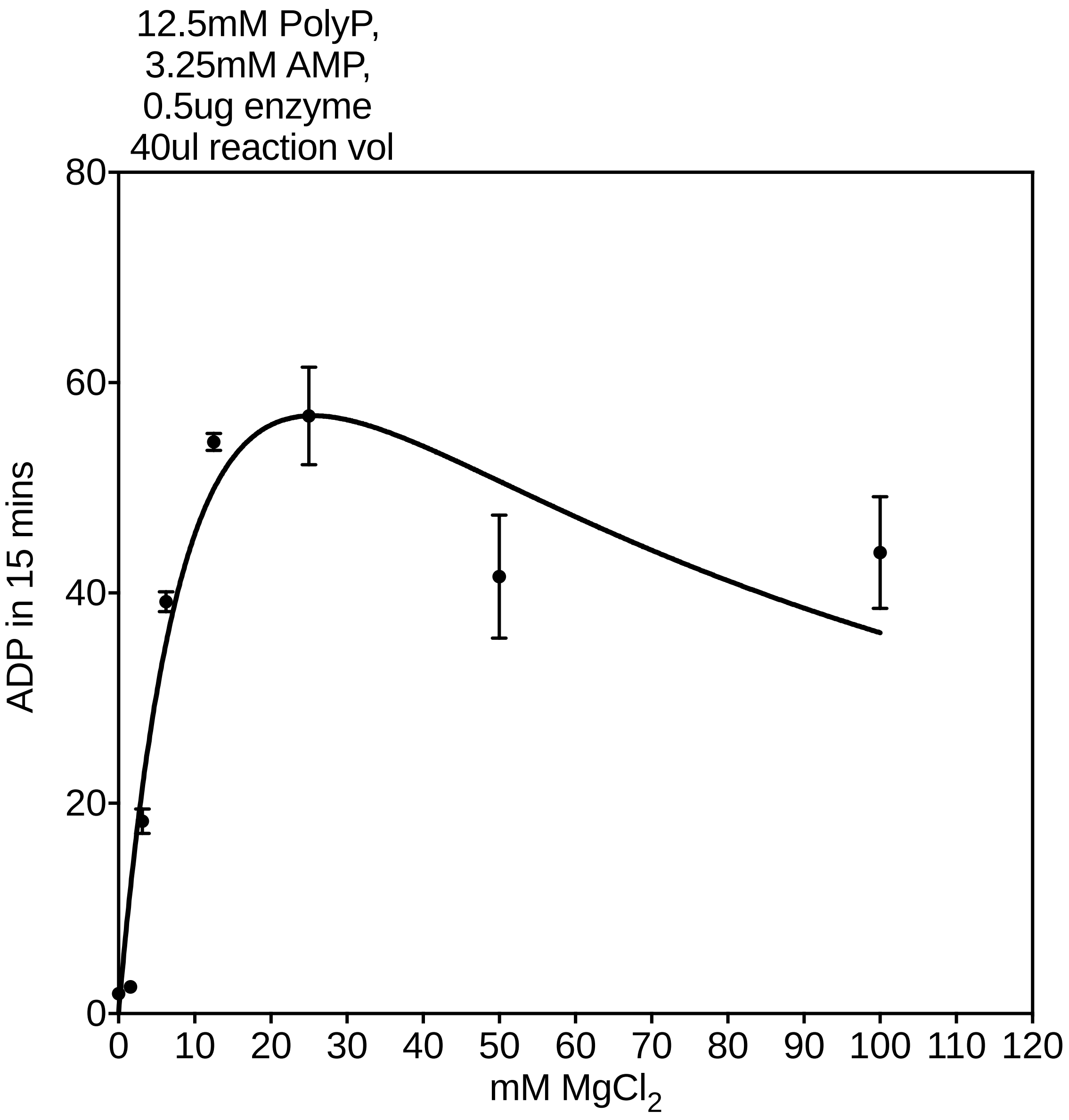


**Supplementary Figure 38 - The kinetics of tnPAP with varying concentrations of $MgCl_2$**
Data have been fitted to the Michaelis-Menten equation with substrate inhibition.

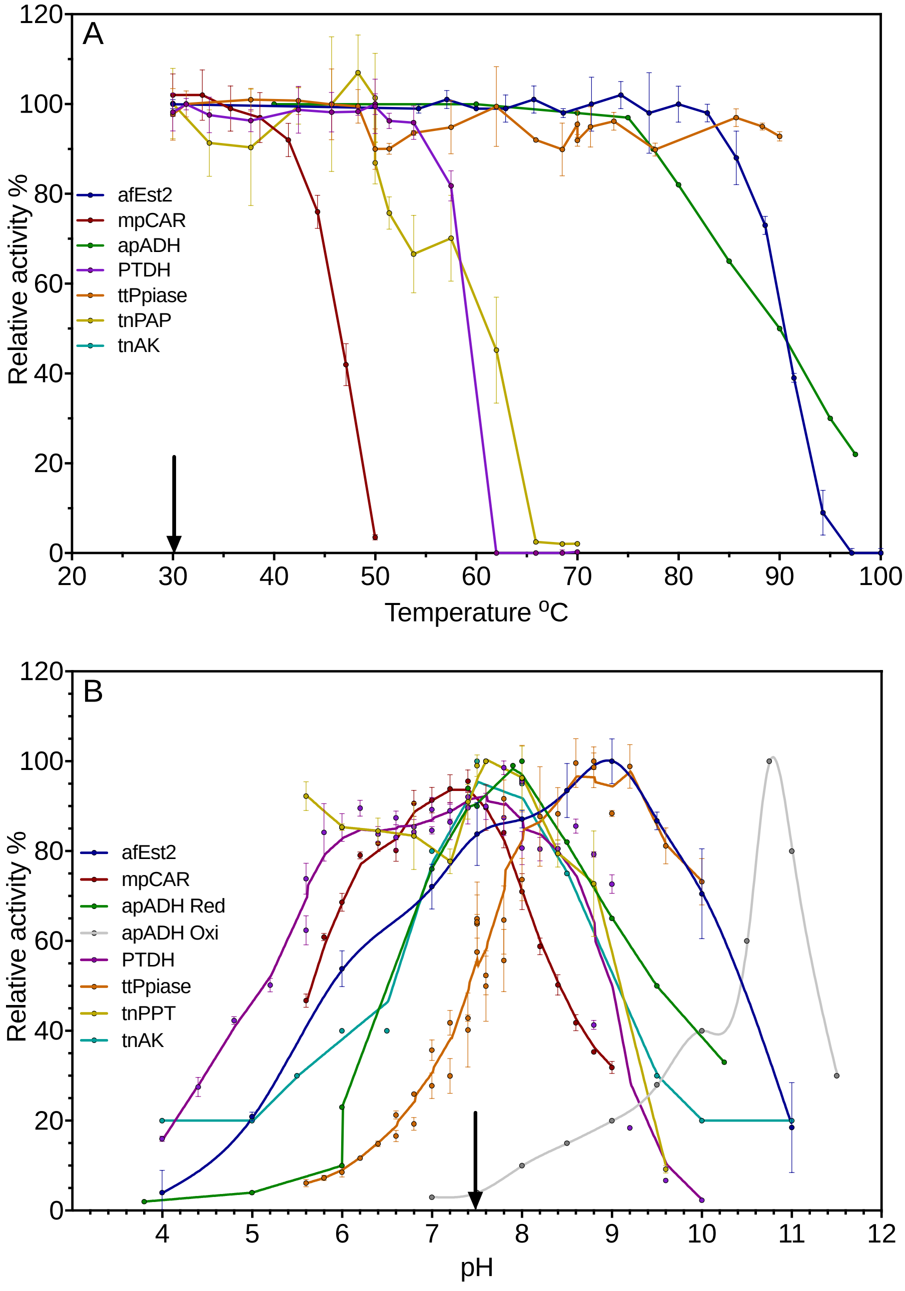


**Supplementary Figure 39 - The operational window for temperature (A) and pH (B) for the seven enzyme reaction**

A. Residual relative activity after incubation at various temperatures for 30 minutes. The selected operational temperature is shown by the black arrow.
B. Relative activity at various pH values. Values are relative to the maximum activity in each case. The selected operational pH is shown by the black arrow.

Data show the mean of three experimental replicates for each point, with error bars representing ±SD. Data for afEst2(42), mpCAR(15), apADH(46) and tnAK (45) were adapted from previous work.

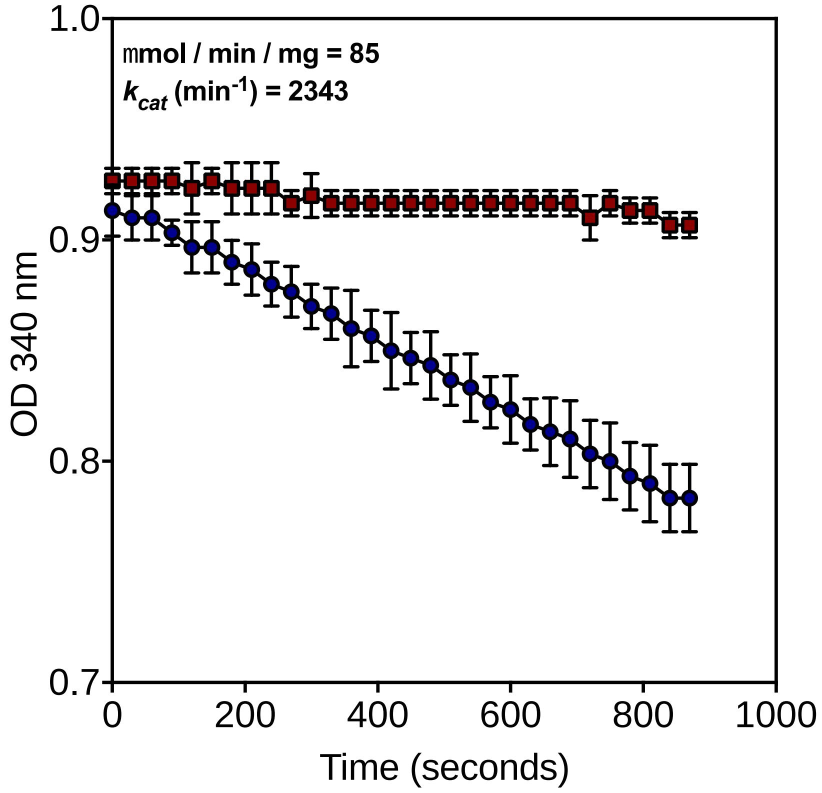


**Supplementary Figure 40 – tnAK – CAR coupled assay to estimate $k_{cat}$**
tnAK was coupled to a CAR enzyme in order to estimate its $k_{cat}$ in the ADP to ATP direction. The rate obtained was significantly slower than previously reported. The $k_{cat}$ for the reverse reaction was adjusted relative. Red circles show the blank rate, blue squares show the rate with tnAK.

**Supplementary Figure 41 – Equilibrium constants for the reactions in the model**
Equilibrium constants were either taken from the literature, or calculated using thermodynamics. Equilibrium constants were used to justify reactions which were modelled as irreversible.

| | **K** | **Modelled as Irreversible or Reversible** | **Ref** |
|---|---|---|---|
| **afEst2** | $4.2 * 10^{-12}$ | Rev | Calculated |
| **PTDH** | $1 * 10^{11}$ | Irr | Woodyer 2003 |
| **Ppiase** | $5.3 * 10^{3}$ | Irr | Davies et al.1993 |
| **CAR** | $7 * 10^{34}$ | Irr | Calculated |
| **PAP** | NA | Rev | NA |
| **AK** | NA | Rev | NA |
| **apADH** | NA | Rev | NA |

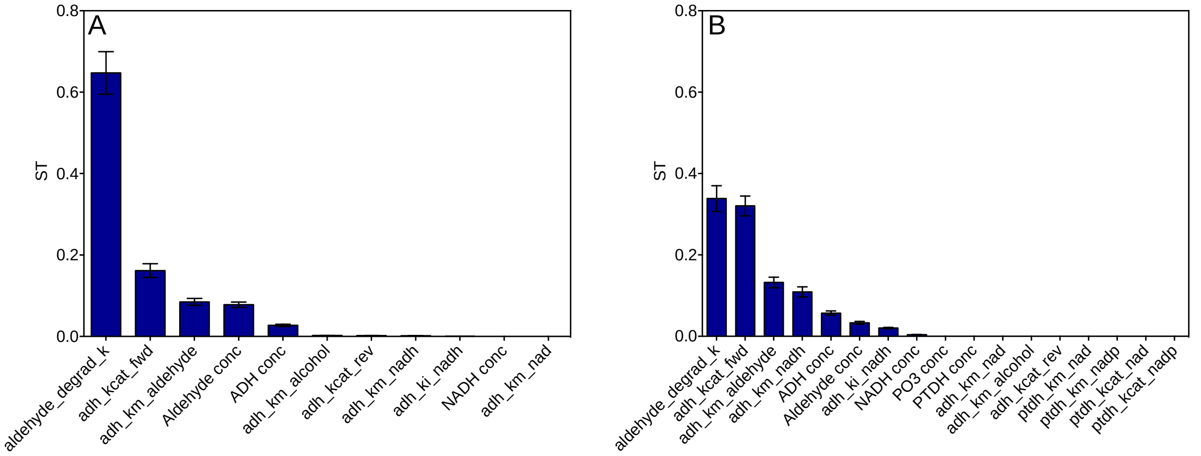


**Supplementary Figure 42 – Sensitivity analysis of apADH and apADH-PTDH modelled reactions.**
The total sensitivity indices (ST) are shown which take into account 1st order and all other interactions. Sensitivity is in reference to the uncertainty in the final *p*-tolyl alcohol concentration. Error bars show the 95 % confidence intervals. The sum of all sensitivity indices' should equal 1.
A: Sensitivity analysis of apADH only reaction, figure 4A.
B: Sensitivity analysis of the apADH-PTDH reaction, figure 4B.

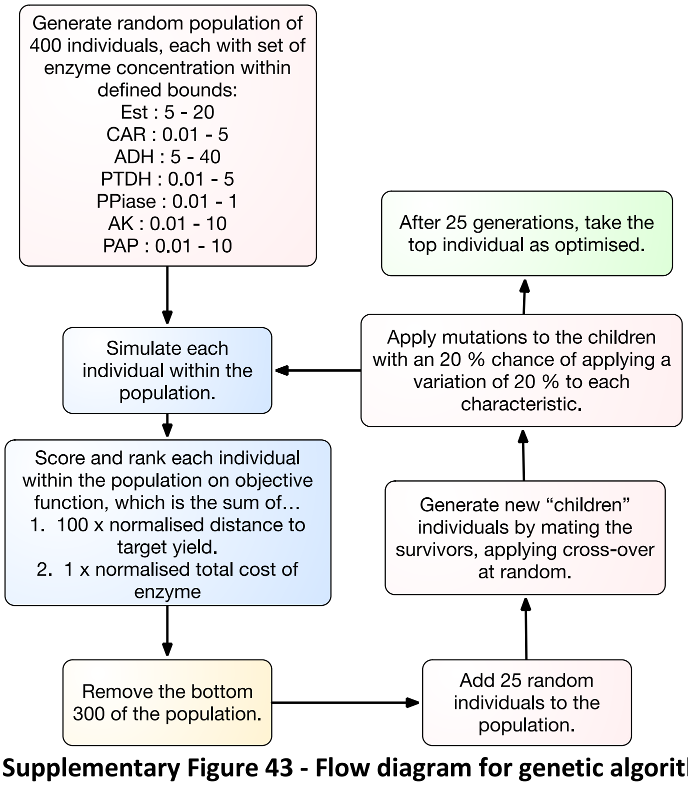


**Supplementary Figure 43 - Flow diagram for genetic algorithm used to optimize the reaction**
Our custom built genetic algorithm was used in the optimization of a batch reaction, minimizing total enzyme cost whilst achieving a target yield of 90 % or above. The flow diagram describes the steps the genetic algorithm carries out to reach this goal.
